# Effects of phenotypic variation on consumer coexistence and prey community structure

**DOI:** 10.1101/2021.06.09.447767

**Authors:** Shane L. Hogle, Iina Hepolehto, Lasse Ruokolainen, Johannes Cairns, Teppo Hiltunen

## Abstract

A popular idea in ecology is that trait variation among individuals from the same species may promote the coexistence of competing species. However, theoretical and empirical tests of this idea have yielded inconsistent findings. We manipulated intraspecific trait diversity in a ciliate competing with a nematode for bacterial prey in experimental microcosms. We found that intraspecific trait variation inverted the original competitive hierarchy to favor the consumer with variable traits, ultimately resulting in competitive exclusion. This competitive outcome was driven by foraging traits (size, speed, and directionality) that increased the ciliate’s fitness ratio and niche overlap with the nematode. The interplay between consumer trait variation and competition resulted in non-additive cascading effects - mediated through prey defense traits - on prey community assembly. Our results suggest that predicting consumer competitive population dynamics and the assembly of prey communities will require understanding the complexities of trait variation within consumer species.

## Introduction

Comparative studies of species traits have historically focused on trait differences between species. However, ecologists increasingly recognize that a large amount of the trait variation in ecological communities exists between individuals from the same species. For example, trait variation within species accounts for up to one-fourth of total trait variation in plant communities[1] and can have ecological consequences comparable in magnitude to those from separate species[2]. Intraspecific trait variation (ITV) may emerge from phenotypic plasticity (e.g., maternal effects or learned behavior)[3], ontogenetic diversity[4], and heritable evolutionary change.

While ITV is ubiquitous in natural ecological communities, its relevance for competitive interactions and species coexistence is unclear. In the past two decades, the relationship between ITV and species coexistence has garnered attention[3, 5] - in particular, the idea that ITV may facilitate the coexistence of competing species[6]. Theoretical studies attempting to link ITV to coexistence have yielded inconsistent results ranging from positive[7, 8], to negative[9, 10], to zero effect[11]. Laboratory and field studies have also demonstrated variable outcomes between competing species with different levels of ITV[12–14]. Currently, this body of empirical and theoretical literature demonstrates no universally positive or negative effect of ITV on coexistence.

Modern coexistence theory is a prevalent theoretical framework in ecology, and it can be leveraged to understand how variable traits (e.g., phenotypic plasticity [3]) might impact fitness or niche differences between competing species. From the perspective of modern coexistence theory, species coexistence or competitive exclusion depends on how a variable trait influences stabilizing and equalizing mechanisms. In empirical studies, the relative contribution of these mechanisms is typically estimated using the niche overlap and competitive (i.e., fitness) ratio computed from population density data fit to a mathematical model of species competition[15]. Still, there are relatively few empirical studies testing current theories of ITV and species coexistence. Furthermore, both theoretical and empirical studies on the effects of ITV and species coexistence have primarily focused on competitive outcomes at a single trophic level. Many studies have shown that ITV of a single consumer can have cascading effects on community dynamics at lower trophic levels[4, 16, 17]. However, little is known about how the interplay between consumer ITV and consumer competition affect prey community dynamics.

In order to uncover general principles of ITV and species coexistence, it will be necessary for empirical studies to better characterize the ecological and evolutionary conditions (including their complexities) that promote or hinder species coexistence. This effort should include understanding the mechanisms (e.g., species traits) that position species’ niche overlap relative to their hierarchical differences and understanding how these operate in broader trophic contexts. Experimental microbial communities offer one way to test these ideas under controlled laboratory conditions. Motivated by these knowledge gaps, we conducted an experiment with a microbial ciliate competing with a nematode worm in a common garden consisting of 24 different bacterial prey species. Specifically, we sought to test 1) how consumer ITV influences the competitive population dynamics between two consumer species, 2) whether links between consumer ITV and interspecific competition influence prey community structure and 3) what mechanisms (i.e., consumer trait differences) might underlie any observed effects.

## Materials and Methods

### Overview

We combined a 24-species isogenic bacterial prey community (each species derived from a single clone, Table S1) with an isogenic nematode worm, *Caenorhabditis elegans*, and an isogenic ciliated protozoan *Tetrahymena thermophila* - hereafter Low Trait Variation (LTV) ciliate. In some treatments, we replaced the LTV ciliate consumer with an even mixture of 20 different ciliate populations, each displaying high intraspecific phenotype variation - hereafter High Trait Variation (HTV) ciliate[18]. In this way, we could manipulate the standing stock of trait diversity in the ciliate consumer in the presence or absence of competition from the nematode. We then combined the LTV ciliate, the HTV ciliate, the nematode, and the prey bacterial community in experimental microcosms with a full-factorial design (six conditions, four biological replicates per treatment). We followed consumer densities, prey density, and bacterial community composition over 61 days representing at least 70 ciliate and 16 nematode generations. We also measured traits from both consumers and prey to determine how community assembly and consumer composition were related to species phenotypes (Fig. 1).

**Figure 1.**
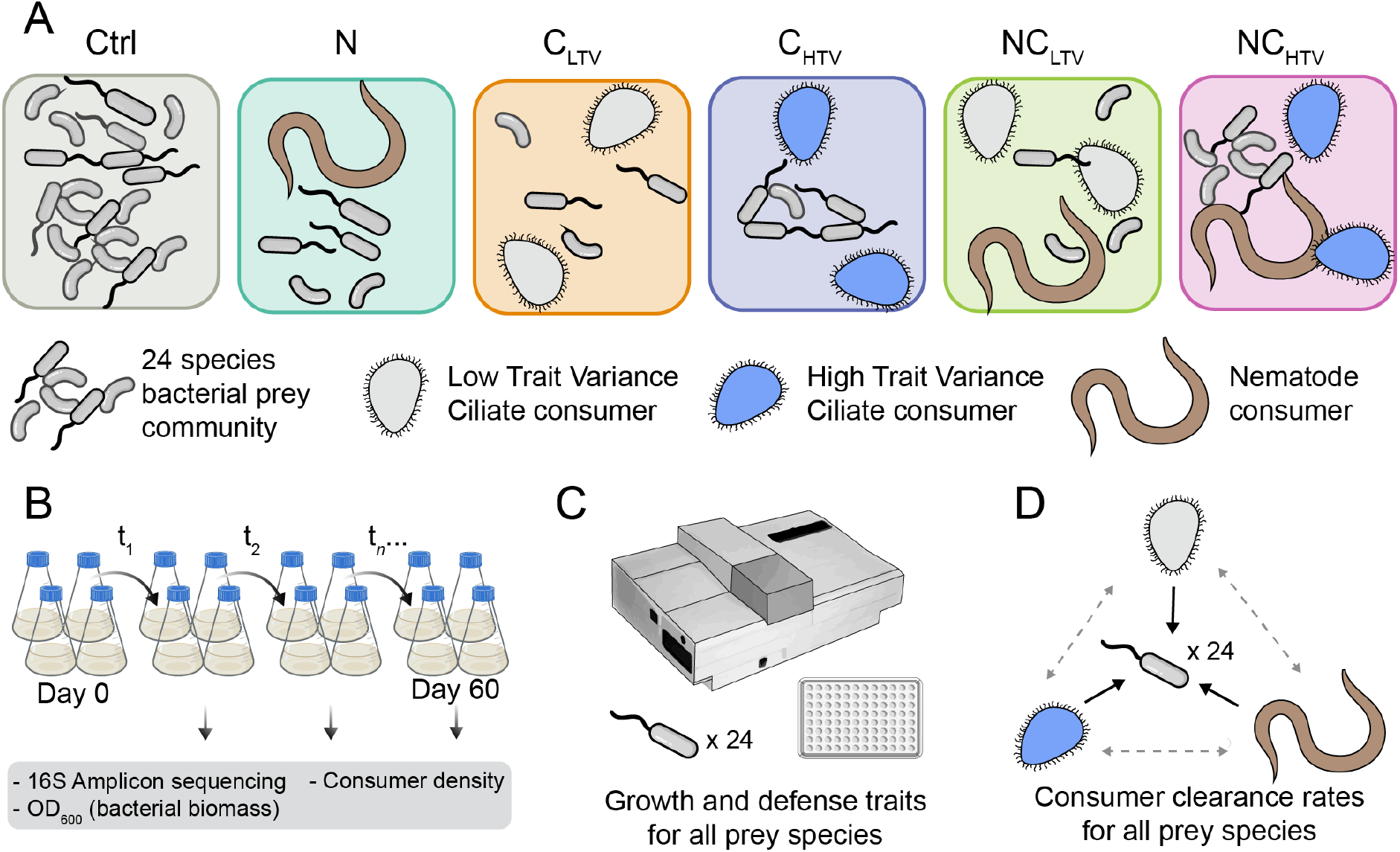
Study setup. **A)** The experimental bacterial community with different consumer combinations. Ctrl - bacteria only, N - bacteria with nematode, C_LTV_ - bacteria with isogenic low trait diversity ciliate, C_HTV_ - bacteria high trait diversity mixed ciliate populations, NC_LTV_ - bacteria + nematode + low trait diversity ciliate, NC_HTV_ - bacteria + nematode + high trait diversity ciliate. **B)** Schematic representation of the serial transfer experiment. Each experimental condition from A) was performed in four biological replicates with 15 transfers/samplings. **C)** Traits of individual bacterial species were measured in monoculture and inferred from genome sequences. **D)** Consumers were grown on each bacterial species to assess consumer grazing efficiency (see methods).

### Study species

The bacterial prey community (Table S1) consisted of 24 phylogenetically diverse species isolated from soil, aquatic, plant, and host-associated environments. Bacteria were grown routinely in 5% King’s Broth and 50% PPY agar. In monoculture, each bacterial species could grow to an optical density of at least 0.3 in the media and temperature conditions used in the main experiment. Each bacterial prey species could support detectable growth of each consumer in pairwise monoculture, but maximum consumer density varied depending on the prey species. The consumers consisted of the ciliate *Tetrahymena thermophila* strain 1630/1U (CCAP) and the clonal hermaphrodite nematode *Caenorhabditis elegans* strain N2. Before the main experiment, *Tetrahymena* was routinely grown by serial transfer in PPY medium[19]. A single *Tetrahymena* mating-type (type II) was used in all experiments and selection lines to ensure that ciliate reproduction was asexual[18, 19]. Nematodes were routinely grown on NGM-plates with *E. coli* OP50 as prey[20]. Revival of consumers and prey from cryostorage was performed as described before[19–21].

### Ciliate intraspecific variation

The mixture of HTV ciliates was prepared by pooling ciliate populations from 20 long-term selection lines (see [18] for details). Briefly, 20 *Tetrahymena* lines were started from isogenic clones (the LTV ciliate progenitor) and continually maintained with one of six bacterial prey species as the sole nutritional source. Four bacterial species were selected to represent genera commonly associated with ciliate predators in soil and aquatic habitats, and two were selected for potential anti-predatory traits. After 600 ciliate generations, we harvested an aliquot from each of these 20 selection lines and made the aliquots axenic using antibiotics. The 20 aliquots were then pooled into a single HTV ciliate treatment. Thus, each HTV *Tetrahymena* inoculum was a heterogeneous population, containing both phenotypic and genetic diversity.

### Experimental setup and sampling

Each consumer treatment in four biological replicates was performed in 20 ml of 5% King’s Broth (KB) liquid medium containing M9 salts and supplemented with cholesterol. Each microcosm was inoculated with 1 × 10^4^ CFU ml^−1^ of each bacterial species. Consumers were inoculated at 1 × 10^4^ ciliate cells ml^−1^ and 10 nematode cells ml^−1^. Microcosms were maintained at 22°C with gentle shaking and transferred to two ml fresh growth medium at day five and then every four days for a total of 15 transfers over 61 days. Nematode and ciliate cell densities were measured directly via microscopy using the Fiji/ImageJ cell counter plugin[22]. Bacterial density was measured using optical density at 600 nm wavelength. All nematodes were counted regardless of size or developmental stage (excluding eggs). Due to methodological constraints, the proportion of individuals in each developmental stage were not formally recorded.

### Trait measurements

Defense against the LTV consumer, carrying capacity, and growth rate for each bacterial strain were measured as described in detail earlier[23]. Maximum growth rate, doubling time, and carrying capacity were estimated using growthcurveR[24]. Biofilm formation capacity was estimated using the crystal-violet method[25]. Bacterial growth was measured on 31 different carbon substrates using BioLog EcoPlates (https://www.biolog.com). All other traits were inferred from bacterial genome sequences using Traitar[26].

To measure pairwise prey clearance by each consumer, each bacterial species was grown as a monoculture in Reasoner’s 2A liquid at 20°C for 96 hours with shaking. Bacterial species were harvested by centrifugation, resuspended in 100 ml of M9 saline solution, then standardized to a common density by adjusting optical density with M9. Five ml of each bacterial species was added to a six-well culture plate with either 2.5 × 10^3^ nematode individuals or 2.5 × 10^4^ of HTV or LTV ciliate individuals. Plates were incubated at room temperature with gentle shaking for 144 hours, after which optical density and consumer counts were conducted. Prey clearance was defined as the difference between the OD_600_ (converted to cells ml^−1^) in each consumer treatment and the no consumer control.

### DNA extraction, sequencing, and analysis

Total DNA was extracted, 16S amplicon sequencing libraries prepared, and sequencing performed as outlined in earlier studies[27]. Read pairs were trimmed and merged into amplicons using BBTools (version 38.61b (https://sourceforge.net/projects/bbmap/). Quality controlled amplicons were assigned to bacterial species using BBMap by mapping against a database of 30 full-length 16S rRNA sequences using the best possible mapping position between the priming sites (see supplementary material for details).

### Statistics and data analysis

All statistical analyses were done in R version 3.6.1[28]. Probabilistic modeling was performed using the Stan programming language v2.24[29] and RStan v2.21.1. We assessed MCMC chain mixing and convergence using effective sample sizes [30] and potential scale reduction factors (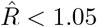 for all covariates)[31]. For all Bayesian analyses, we used default or weakly informative priors.

Ciliate, nematode, and bacterial densities were modeled with hierarchical generalized additive models using mgcv v1.8-33[32]. All other regression was performed in the Bayesian framework using rstanarm v2.21-1 [33] or MCPv0.3-0[34]. Differences in consumer competitive ability and niche overlap were estimated by parameterizing phenomenological Lotka-Volterra competition models. The inferred per capita growth rates, intraspecific, and interspecific coefficients were then used to calculate competitive and niche differences[15]. The compositional bias of amplicon-derived species abundances was estimated using metacal v0.1.0[35] and abundance information from control samples (day 0) where all prey species were inoculated at equal cell densities (1 × 10^4^ CFU ml^−1^). The Shannon diversity dissimilarity index was calculated following Jasinska *et al.*[36]. Population-level alpha and beta diversities (Shannon diversity and Bray Curtis dissimilarity) were estimated using DivNet v0.3.6 [37]. Segmented regressions of Shannon diversity were performed using MCP v0.3-0[34] to identity sorting and equilibrium phases. Beta diversity for individual replicates was calculated using Bray and Curtis dissimilarity on species relative abundances and non-metric multidimensional scaling for ordination.

The HMSC package[38] was used for joint species distribution modeling. A two-step hurdle model approach was used to account for zero inflation in the log-transformed count data where first a probit model for presence/absence was fit and then a Gaussian model for regularized log-transformed species abundances conditional upon presence. Differences in sequencing depth between libraries were controlled for by including log-transformed total reads as a fixed effect in the model. Sequencing count data is compositional and statistical approaches for absolute abundances should be applied cautiously due to a risk of artifacts. However, we also emphasize that bacterial population density (as measured by optical density) was not significantly different between predation treatments (see results), suggesting that the compositional differences are likely to reflect true differences. The hurdle model approach is commonly used for modeling sequencing count data with HMSC[38–40]. Details about the model formulation and testing are described in the supplementary material.

## Results

### Consumer ITV determines competitive exclusion

The magnitude of ITV in the ciliate consumer was sufficient to alter the ecological dynamics of the competing consumer. Competition between the LTV ciliate and the nematode excluded the ciliate by day 11, while competition between the HTV ciliate and the nematode excluded the nematode after 40 days (Fig. 2A). These changes were not due to altered consumer or prey carrying capacities due to ciliate ITV. In the absence of competition, the carrying capacity of the HTV ciliate was not different from the LTV ciliate growing on the prey community (Table S2). Prey biomass differed by a mean fold difference of 1.08 across all consumer treatments and sampling times (Table S3), indicating that each consumer or consumer pair drew down total prey resources to similar levels. Therefore, the population dynamics of the LTV and HTV ciliate in monoculture were effectively equivalent.

**Figure 2.**
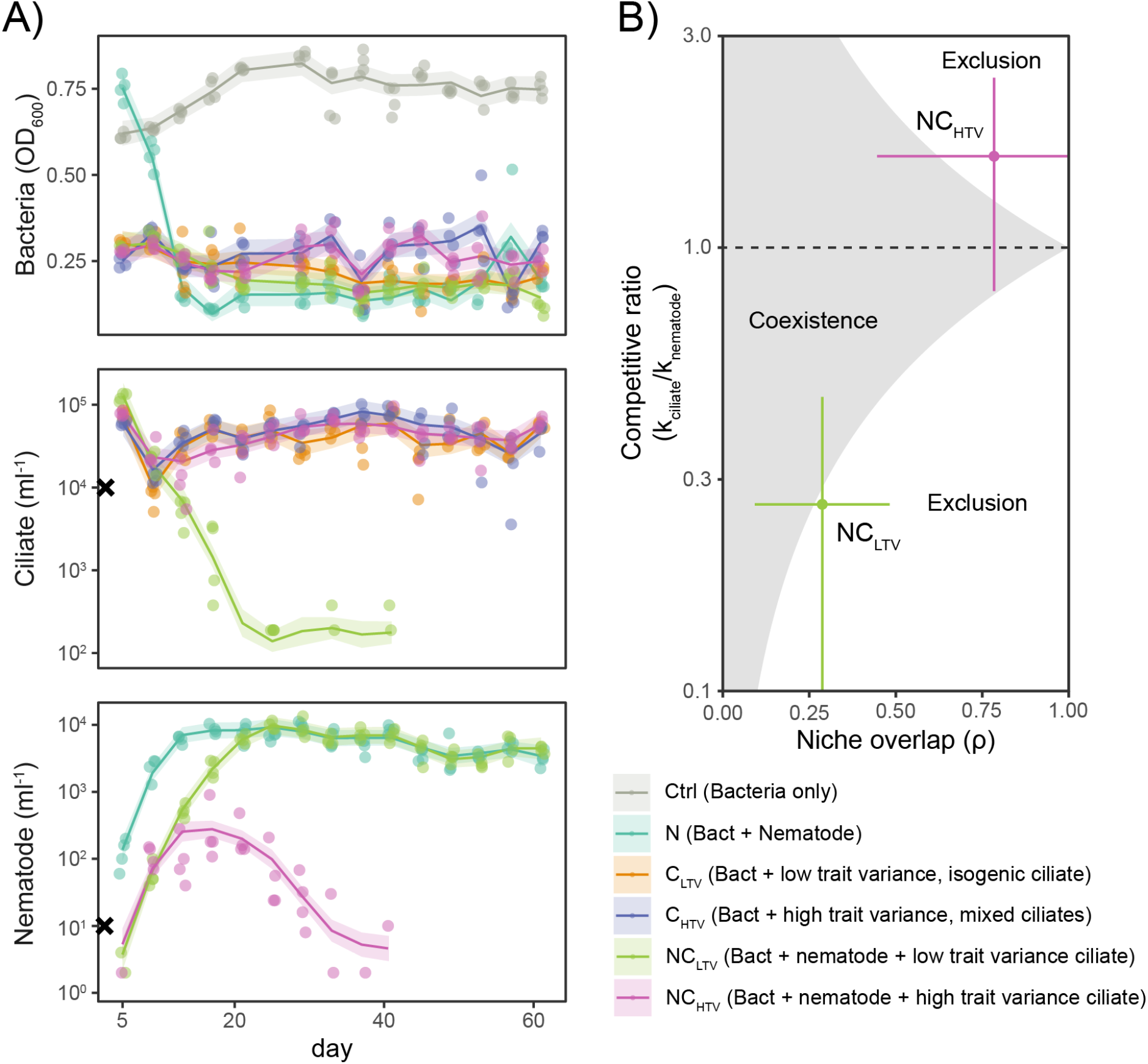
Consumer competitive hierarchy depends upon ciliate ITV. **A)** Consumer and bacterial biomass during the course of the experiment. Points are observations, lines are estimated marginal means (mean response and 95% confidence levels) from generalized additive models. Black crosses are target consumer starting densities at T_0_. Model summaries and contrasts of marginal means are available in Tables S2-S4). **B)** Niche overlap and competitive ratio between the ciliate and the nematode depending on whether the ciliate had low trait variation (NC_LTV_) or high trait variation (NC_HTV_). Coexistence is defined by the inequality 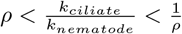. Points and line ranges shows the mean and standard deviation from four biological replicates.

We next attributed consumer dynamics to changes in niche overlap and their competitive ratio following coexistence theory. Coexistence theory describes the coexistence of two species as a balance between stabilizing and equalizing mechanisms[41]. Equalizing mechanisms relate to inherent competitive differences between species that describe dominance hierarchies in a community. Stabilizing mechanisms (i.e., niche *differences*) relate to the ability of each species to recover from low density after a perturbation. ITV could potentially impact these two mechanisms in any number of ways. We estimated niche overlap and the competitive ratio of the HTV or LTV ciliate to the nematode by parameterizing the competitive Lotka-Volterra equations and obtaining the resulting interaction coefficients (see methods). This approach is phenomenological because it represents the outcome of competition/coexistence rather than an underlying biological mechanism per se. Nevertheless, it can be informative for understanding the balance of forces acting upon competing species in natural communities.

Relatively strong niche differences separated the LTV ciliate and nematode, but a strong competitive advantage for the nematode exceeded these differences (Fig. 2B). The model predicted that the LTV ciliate and nematode were at or near the boundary of coexistence. In both HTV and LTV competitive arrangements we observed very low abundances of the inferior competitor (*<* 100 ciliates ml^−1^, < 5 nematodes ml^−1^) in some of the replicates near day 40. Accurate manual counting is challenging at low densities, and it may be that the inferior competitor never truly became extinct. Finally, ciliate ITV strongly increased the ciliate/nematode competitive ratio while simultaneously increasing niche overlap. The magnitude of the competitive change was much larger than the niche change, which suggests that ciliate ITV predominantly drove hierarchical competitive differences between consumers (Fig. 2B).

### Consumers drive prey community assembly

We next sought to determine the degree to which prey communities were structured by consumer treatment versus stochastic processes (e.g., variability in composition or density of prey inocula or random birth and death processes). We expected phenotypic heterogeneity in the HTV ciliate might increase the likelihood of early stochastic events driving divergence in prey community composition[42]. We surveyed the repeatability of prey communities across biological replicates within treatment categories by calculating the scaled Shannon diversity dissimilarity (*D′*) across treatments at each sampling point. The observed *D′* values (Fig. S1B) were close to 0 (highly similar replicates) and significantly lower than expected if cross-replicate variance equaled cross-treatment variance (Table S5). Surprisingly, the HTV ciliate generated equally repeatable prey assembly compared to the LTV ciliate or the nematode. These results show that 1) technical variation in laboratory conditions (e.g., temperature or transfer volumes) were minimal for these experiments, and 2) the process of prey community assembly alongside each consumer or consumer pair was strongly deterministic.

Consumers significantly altered prey species’ local abundance and evenness (i.e., alpha diversity). Prey communities generally assembled under each treatment so that the minority of sequencing reads represented the majority of species. (Fig. 3A). Most communities were numerically dominated by nine prey species (Fig. S1). Shannon diversity varied in two distinct phases across treatments, which we defined operationally (Fig. S2, supplementary methods). In the first phase (hereafter sorting), Shannon diversity declined linearly with time, while in the second phase (hereafter equilibrium), it oscillated around a temporally zero-trend mean (Fig. 3B). The presence of a consumer or co-consumer pair slowed the rate of Shannon diversity decline in the sorting phase relative to the control (Fig. S3, Table S6). In the equilibrium phase, predation also generally supported higher mean Shannon diversity. However, the HTV ciliate generated significantly higher prey diversity than the LTV counterpart (Table S6). These findings are consistent with the expectation that interspecific competition between prey is reduced in the presence of predation [43]. Like prior studies in other systems[4, 16, 17] we find evidence that ITV has cascading effects on prey communities - here resulting in increased prey diversity. However, consumer competition did not change either the rate of diversity decline in the sorting phase or the mean diversity in the equilibrium phase relative to consumer monocultures (Tables S6-S7). Instead, the effect of two competing consumers on the Shannon diversity change was generally equivalent to that of a single consumer.

**Figure 3.**
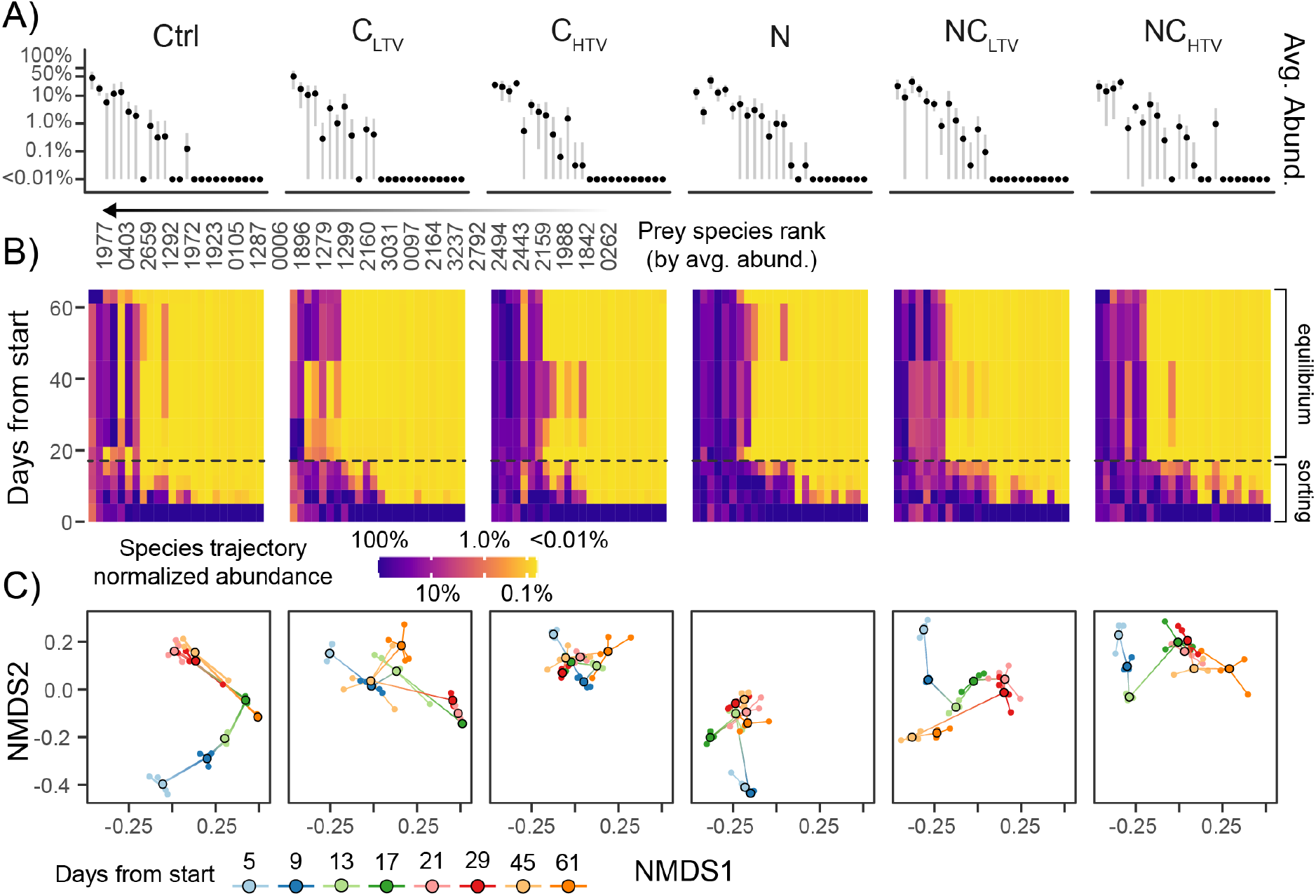
Prey community response to consumer competition and ITV. **A)** Prey species abundance distributions from each experimental treatment. X-axis ranks prey species (1977 most abundant) by average across all conditions. Y-axis shows relative abundance. **B)** Replicate mean abundance trajectories over time (normalized to each species maximum) for each of the 24 bacterial prey species. **C)** Non-metric multidimensional scaling ordination of trajectories through community space. Larger, outlined circles show means across replicates, while small circles are replicates. Colors represent days from the experiment start. Means are connected in chronological order.

We then examined the separation of prey communities across time and consumer treatments (i.e., beta diversity) using ordination to visualize the trajectory of each community through system phase space. The distance between ordination points can be interpreted as a measure of system stability where larger “jumps” through state space represent periods of relative flux[44]. Overall, ciliate ITV resulted in prey communities that were more stable through time than prey communities forming with the LTV ciliate (Table S8). The LTV ciliate and no-consumer prey communities traversed the longest distance in community space, indicating these communities were the least stable through time (Table S8). However, prey communities assembling with the HTV ciliate and nematode showed relatively small changes in beta diversity and stabilized rapidly (Fig. 3C). Prey communities forming alongside competing consumers were compositionally distinct from those forming with either consumer by itself. The presence of two competing consumers in the first 20 days generally had a destabilizing effect on prey temporal trajectories (larger jump lengths) relative to single consumer treatments. Thus, consumer ITV provided a stabilizing effect on prey communities while consumer competition was generally destabilizing.

### Links between consumer ITV and competition have non-additive effects on prey abundances

We next asked how consumer ITV and competition influenced community-level properties of the bacterial prey. We used a joint species distribution model (JSDM) [45] to simultaneously model the responses of the 24 bacterial prey species to different consumer competitive arrangements while estimating the contribution of traits and phylogenetic relatedness to the prey community response (see methods). We modeled the sorting and equilibrium phases separately to account for the strong temporal dependence in the sorting phase (supplementary text). The average explanatory power was excellent for the sorting phase model (*R*^2^ = 0.90 ± 0.09) and good for the equilibrium phase model(*R*^2^ = 0.58 ± 0.19), both of which showed the most predictive power through their fixed effects (Table S9). Variance partitioning over the explanatory variables showed that model fixed effects explained a substantial amount of variance in species abundance (Fig. 4). These findings demonstrate that the JSDMs sufficiently described the observed prey abundance data.

**Figure 4.**
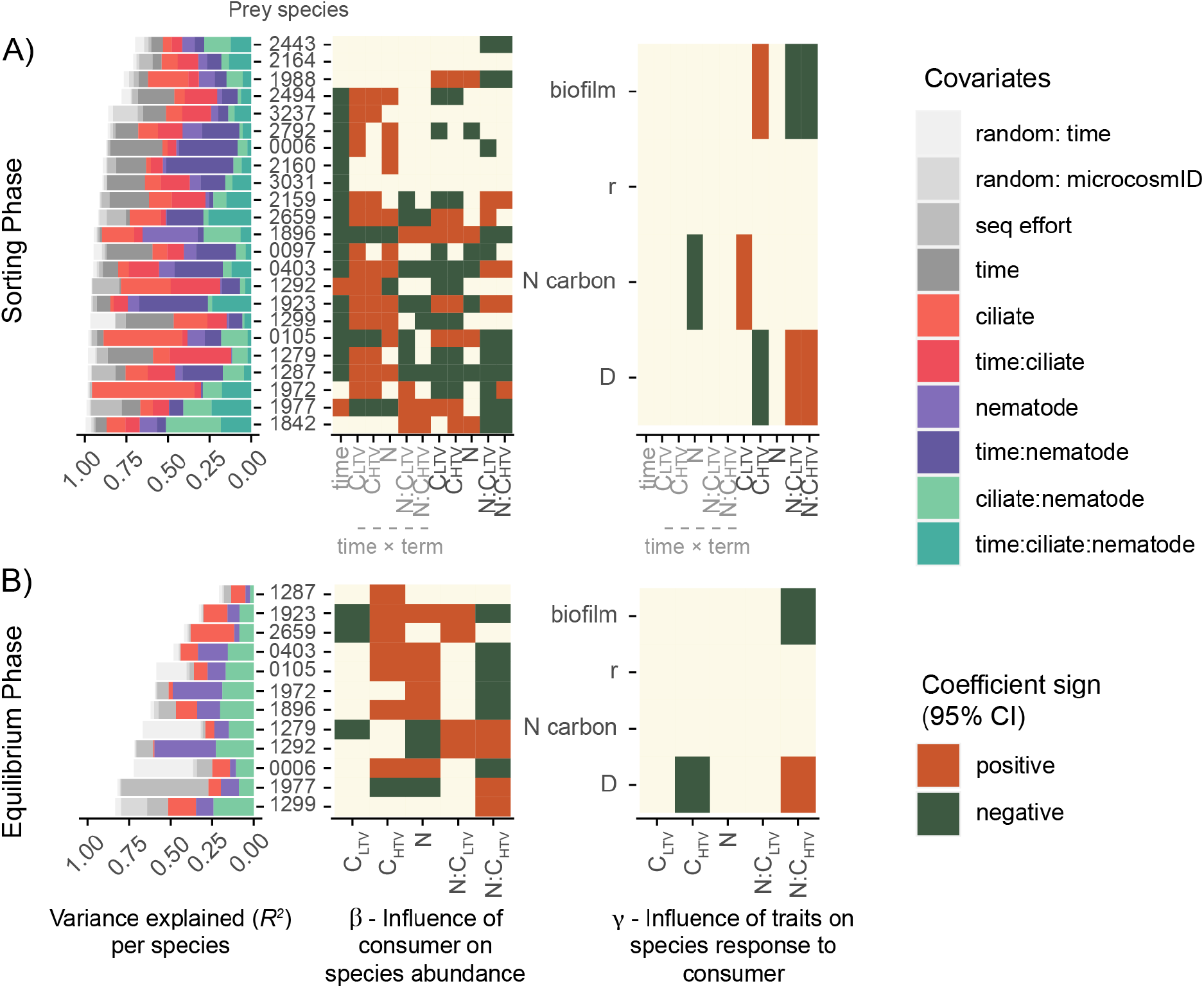
Determinants of prey community assembly. JSDM results from **A)** the sorting phase (days 0-13) and **B)** the equilibrium phase (days 17-61). Bar plots show the explanatory power for each bacterial prey species in the JSDMs. The bars are colored by the proportion of variance explained by fixed and random effects and their interactions. Heatmaps show the influence of fixed effects on bacterial prey species abundance (β) and the influence of prey traits on bacterial prey species association with fixed effects (γ). The sign of the coefficient (i.e the direction of the effect) is included only if the 95% credibility interval excludes zero (i.e the probability the effect is not zero is 95%). Trait abbreviations: D, defense against C_LTV_; N carbon, number of carbon sources the species can use; r, maximum specific growth rate; relative biofilm, biofilm production.

Consistent with the beta diversity patterns we observed above, we found that consumer competition resulted in significant non-additive effects on the abundances of both common and rare prey species. Specifically, prey composition with two consumers was distinct from communities with either consumer alone. We define non-additive effects as the presence of a non-zero coefficient for an interaction term in the JSDM. In practice, this means that prey species were significantly less or more abundant than predicted by a linear combination of the individual predator effects (i.e., the effect of one consumer depended on the presence of the other). In the sorting phase, time had a negative effect for most species, reflecting the rapid decline of rare species from their initial frequencies (Fig. 4). The consumer-specific changes in time were largely positive, indicating that most species increased over time relative to the no-consumer control. Over 60% of prey species in the sorting phase had a non-zero coefficient for the LTV ciliate-nematode (C_LTV_ × N) and HTV ciliate-nematode (C_HTV_ × N) interactions. These two interaction terms were of opposite sign to the individual consumer effects, but they were usually in the same direction. This implies that non-additivity was primarily driven by the presence of two consumers and not consumer ITV. The nematode effect had the greatest explanatory power in the equilibrium phase, followed by the ciliate-nematode interaction term. In contrast to the sorting phase, the consumer interaction effects were concentrated in the C_HTV_ *×* N interaction, while the effect of the LTV ciliate and nematode were additive (95% of the posterior distribution of the C_LTV_ × N interaction term overlapped with zero). Overall, ITV strengthened the non-additivity of multiple consumers on prey abundances in the experiment’s equilibrium phase.

### Prey community responses are mediated through defense traits

We next used the JSDM framework to test whether phylogenetic relationships and prey traits were associated with specific community responses to the different consumer arrangements. We included traits for defense, growth rate, biofilm formation, and the number of carbon compounds degraded and used by each prey species. Defense, growth rate, and biofilm formation are likely to be important traits for consumer-prey interactions. We used the number of carbon compounds as a generic proxy for catabolic versatility, which we expected might be relevant for both competitive and cooperative interactions between prey species. Collectively, these four traits represented approximately 80% of the variance from the total collection of all species traits we surveyed (Fig. S4). The 95% credibility interval of the phylogenetic posterior included zero and the median effect was small (sorting=0.05 × 0.11, equilibrium=0.32 ± 0.34). Thus, closely related bacteria were no more likely to share a common response to the consumer treatments than distant relatives. Prey traits explained a substantial amount of variation (22% and 44%) in both experimental phases (Table S10, Fig. 4), suggesting a shared phenotypic response of the prey community to each treatment.

The influence of prey traits on prey community assembly with competing consumers was also generally non-additive (Fig. 4). Prey traits explained the greatest proportion of the community response to the HTV ciliate and the C_HTV_ × N interaction (Table S10). For example, prey traits explained up to 80% of the variance of species response to the C_HTV_ × N term. In particular, prey defense and biofilm traits were consistently associated with community responses to the HTV ciliate either alone or through an interaction with the nematode in both experiment phases (Fig. 4). This finding suggests that the non-additive response of prey species to consumer competition and ITV were mainly attributable to differences in prey traits.

### Effects of ITV on the consumer response

In the short term, a consumer with uniform clearance rates across prey species should remove those prey species in proportion to their abundance in a consumer-free community. Alternatively, a consumer displaying nonuniform prey clearance (i.e., prey preference) should remove prey in a way that deviates from this proportionality[46]. We asked how consumer ITV altered the overall consumer response by estimating foraging preferences from sequencing data of each consumer and the prey community. We regressed the relative abundance of each bacterial species in the presence and absence of each consumer across all samples from the sorting phase of the experiment. Slopes approaching one indicate uniform prey removal, while slopes approaching zero indicate strong consumer preference[46]. We found that the slope was largest for the LTV ciliate (Fig. 5A), suggesting that it had the smallest relative variation in clearance across all prey species. Both the HTV ciliate and nematode slopes were significantly smaller than the LTV ciliate slope, indicating that both these consumers preferentially foraged on some prey species (Table S12) - particularly highly abundant prey.

**Figure 5.**
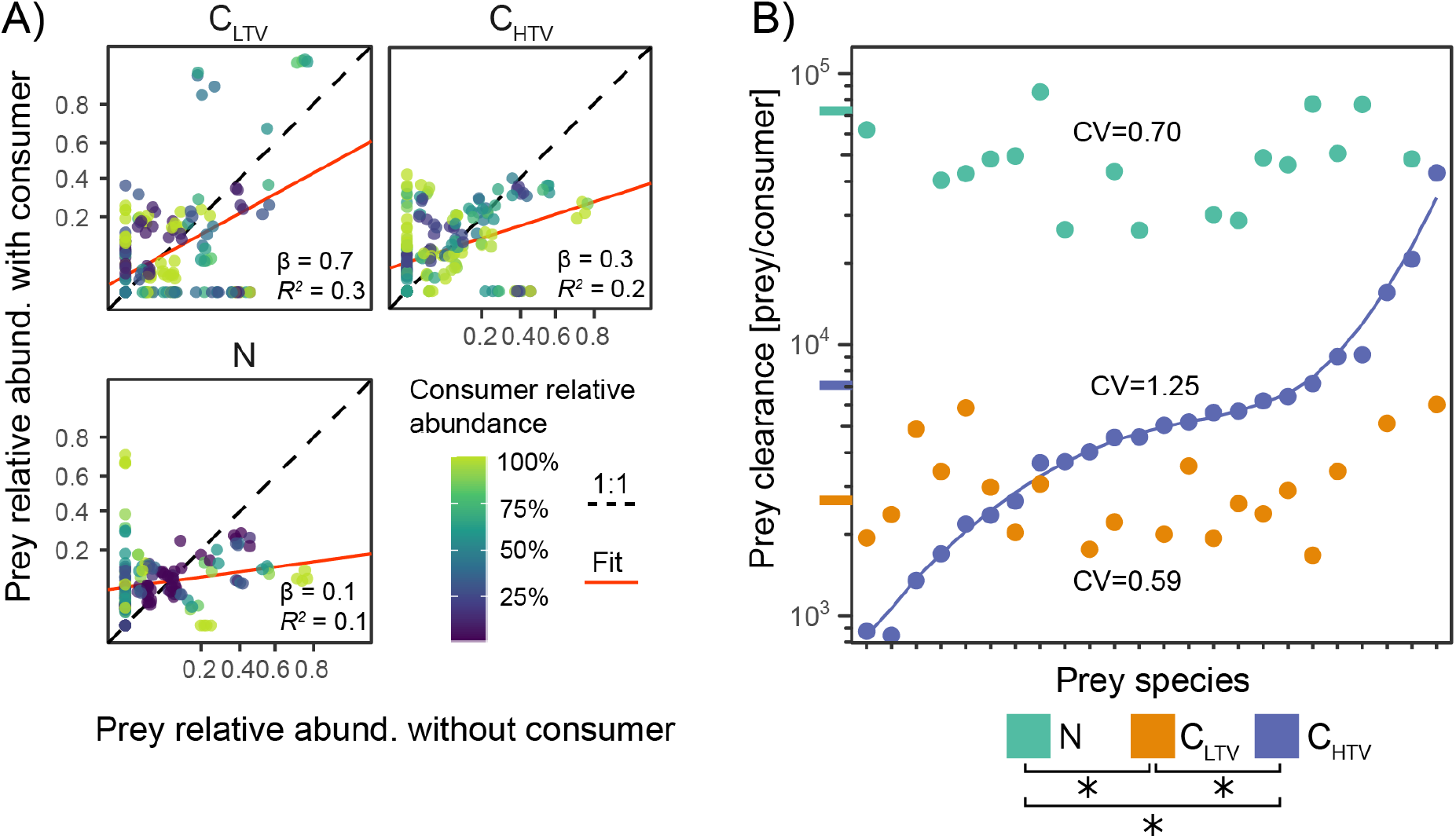
Consumer ITV drives partitioning of the predation response. Relative abundance of prey species in the presence of ciliate or nematode consumers (vertical axis) versus the corresponding relative abundance in the consumer-free controls. The regression includes prey species > 1% relative abundance and samples from first 21 days. The axes are arcsin square-root transformed. *β*, *R*^2^ is the median for the posterior of the slope and Bayesian *R*^2^, respectively from linear regression (Table S12). B) Consumer response (prey clearance) as a function of prey species identity. Prey clearance is the number of bacterial cells removed per consumer individual after 144 hours. Long tick marks on the vertical axis show the mean clearance rate of each consumer. Prey species are ordered by increasing susceptibility to the HTV ciliate. The curve is a simple polynomial fit to aid visualization. Asterisks show significant differences in mean prey clearance between consumers (Table S11). CV = coefficient of variation.

We next tested whether the nonuniform foraging we inferred from the sequencing data matched experimental observation. We experimentally assayed the ability of each consumer to clear prey biomass in pairwise grazing trials with each of the 24 bacterial strains (Fig. 5B). The results were consistent with our expectations from the sequencing data: 1) the LTV ciliate removed the fewest prey cells per capita with the most uniform clearance rates (smallest coefficient of variation) across prey species, 2) the HTV ciliate feeding performance increased for some prey species but decreased for others (resulting in a large coefficient of variation), and 3) the nematode removed the most prey per capita but with a similar uniformity across prey species to the LTV ciliate. Overall, ITV increased mean ciliate grazing performance as well as its capacity for selective foraging.

## Discussion

Theoretical and empirical studies have documented ITV both promoting and undermining the coexistence of competing species[7–10, 12–14]. Modern coexistence theory could potentially reconcile many of these conflicting outcomes as it offers a highly general framework for compartmentalizing the species parameters affecting coexistence into equalizing and stabilizing mechanisms[41]. To successfully understand ITV and species coexistence using modern coexistence theory, we require knowledge of the varying traits and intuition into how they map to competing species’ competitive ratio and niche overlap. High ITV in traits affecting niche overlap (niche traits) should result in a distribution of individuals both adapted and maladapted to the prevailing environmental conditions. We expect that ITV in niche traits will weaken stabilizing mechanisms because conspecifics can distribute over a wider niche breadth, which increases niche overlap with heterospecifics. Alternatively, high ITV in traits altering species fitness ratios (competitive traits) would make some individuals more fit and others less fit under a given environmental condition. The predicted effects of ITV on competitive traits should depend upon the shape of the trait-response curve due to Jensen’s Inequality[9, 47]. Therefore, the expectation for coexistence will also depend upon the shape of the species performance response to the variable trait. Furthermore, the fitness effects of ITV on competitive traits will depend upon the mean trait value and the magnitude of variance - for example, whether the average individual is adapted or maladapted to the local conditions [48]. Traits may also simultaneously impact both niche and fitness differences in complicated ways[49].

We previously reported that ciliate individuals from the HTV populations were, on average, 40 *μ*m larger, 1.25× faster, and turned less often than isogenic, LTV ciliate individuals[18]. Individuals with high values of these traits should cover a larger search volume than smaller, slower individuals[50]. Assuming a spatially homogeneous distribution of prey cells, this should allow HTV ciliates with extreme trait values to encounter more prey per unit time than the average LTV individual. Here we expect that being larger, faster, and swimming in straighter trajectories would affect *both* the fitness ratio and niche overlap between the HTV ciliate and the nematode. Nonetheless, we propose that these traits are primarily competitive. This idea is consistent with the increased HTV ciliate prey clearance averaged across all prey species (Fig. 5A) and the large increase in the competitive ratio between the ciliate and nematode (Fig. 2B). However, we also suspect that ITV in these competitive traits weakened stabilizing mechanisms. Most of the HTV ciliate fitness gains were concentrated on prey species already preferred by the nematode, which was reflected in the prey clearance trials (Fig. 5) and the estimates of niche overlap from the competitive Lotka-Volterra model (Fig. 2). Overall, our findings are consistent with predictions from modern coexistence theory, which broadly suggest that ITV should hinder species coexistence[9, 10]. However, these studies also suggest that ITV might promote coexistence if it allows a mean-variance trade-off favoring the weaker competitor[9, 10, 51]. We found that ITV in the weaker competitor also hindered coexistence because there was no generalist specialist trade-off[10]. In general, the HTV ciliate was a more effective consumer than the LTV ciliate, and the HTV performance gains disproportionately impacted the nematode’s optimal prey. Thus, the competitive differences between the two consumers were also reinforced by the convergence of their niches.

The link between consumer ITV and competition also strongly influenced the community composition of the prey trophic level. The HTV ciliate or the nematode rapidly stabilized the composition of the prey community, while the LTV ciliate and no-consumer treatment caused the prey community to make the largest oscillations through community state space (Fig. 3C). The combination of two consumers caused the prey community to traverse state-space distinctly from either consumer alone. Indeed, we found that the combined effects of the ciliate and nematode on individual prey species abundances were often non-additive and that these non-additive effects were mediated through prey species defense traits (Fig. 4). Consumer trait diversity generally strengthened the non-additivity of how prey defense traits influenced prey community assembly in the presence of the competing consumer. It is important to note that the non-additive prey responses may not necessarily be a direct consequence of predation (e.g., interference between predator species or altered prey behavior in response to predation) but rather a result of predation-driven shifts in community composition that promoted alternative higher-order interactions between prey species (i.e., crossfeeding) or cascading effects due to prey extinctions[52]. It is also possible that consumer feedback loops result in apparent mutualistic or competitive outcomes for prey in the absence of direct consumer-prey interaction[53]. Previous studies have also shown that consumer ITV has cascading effects on the composition of prey communities and ecosystem function [4, 16, 17]. Here we show that consumer competition and consumer ITV interact to influence the assembly of bacterial prey communities.

There are some caveats to consider when translating our findings to other ecological systems. First, we do not know whether the experimental outcomes with the HTV ciliate were due to the dominance of a single ciliate genotype or a combination of strains. However, aggregate phenotype data from the HTV populations suggest that the magnitude of change in size, speed, and directionality traits was relatively modest compared to the LTV ciliate[18]. Thus, we assume that the outcomes with the HTV ciliate were a consequence of the total diversity in the population and not the superiority of a single genotype/phenotype. Nevertheless, we cannot exclude the possibility that some HTV ciliate genotypes had more impact than others. Future studies may be able to quantify the competitive effects of genotype on niche and competitive traits. Second, we did not explicitly manipulate the amount of ITV in the nematode competitor, although phenotypic variation was almost certainly present as developmental diversity or potentially maternal effects. Although we did not formally record *C. elegans* developmental stages, the size distribution was uniform across treatments and sampling times. Thus, we expect any indirect effects from the developmental structure in the nematode populations were primarily shared between the HTV and LTV ciliate treatments. Still, we cannot completely exclude the possibility that nematode ontogenetic structure might play a relevant role in the competitive dynamics we observed.

Although a fast-paced body of theoretical work addresses ITV and species coexistence through modern coexistence theory, there is not yet a unified framework reconciling the various predicted outcomes. Empirical work can contribute to this challenge by testing theoretical predictions. Our results are broadly consistent with the predicted effects of ITV on species coexistence from the modern coexistence theory framework. However, our findings also highlight some important mechanistic details. We found that variable traits in the ciliate predominantly functioned as competitive traits, but they also impacted niche overlap with the nematode by partitioning its response to the nematode’s preferred prey species. Our intuition is that many traits may act simultaneously as equalizing and stabilizing mechanisms, as we observed here. Alternatively, traits may also influence niche overlap or fitness ratios differently depending on ecological context. Moving forward, it will be necessary to better understand such complexities and how this affects species coexistence more generally.

## Data availability

Raw sequencing data is available from the NCBI Sequence Read Archive (SRA) under the BioProject accession number PRJNA725120. Preprocessed count tables are available from https://github.com/slhogle/consumer-competition.

## Code availability

Scripts reproducing all figures and steps in the data analysis are available from https://github.com/slhogle/consumer-competition.

## Acknowledgments

Elizaveta Zakharova: investigation - supporting, data curation. Olli Pitkänen: investigation - supporting, data curation. Ida-Marija Hyvönen investigation - supporting, data curation. SLH thanks Chuliang Song for helpful discussion. TH dedicates this work to the memory of Dr. Jouni Laakso. Funding: Academy of Finland (grants #327741 to TH and #286405 to LR)

## Author contributions

SLH: conceptualization, supervision, software, formal analysis, writing - original draft. IH: conceptualization, investigation - lead, visualization, writing - review & editing. LR: supervision, formal analysis, funding acquisition, writing - review & editing. JC: conceptualization, writing - review & editing. TH: conceptualization, supervision - lead, project administration, funding acquisition, writing - review & editing.

