## Supplementary material for "Effects of phenotypic variation on consumer coexistence and prey community structure"

**Author Contributions:** SLH: conceptualization, supervision, software, formal analysis, writing - original draft. IH: conceptualization, investigation - lead, visualization, writing - review & editing. LR: supervision, formal analysis, funding acquisition, writing - review & editing. JC: conceptualization, writing - review & editing. TH: conceptualization, supervision - lead, project administration, funding acquisition, writing - review & editing. All authors provide approval for publication.

**Competing Interest Statement:** The authors declare no conflict of interest.

**Preprint Servers:** The article is available on the bioRxiv preprint server: `https://www.biorxiv.org/content/10.1101/2021.06.09.447767v1`.

**Data Availability:** Raw sequencing data is available from the NCBI Sequence Read Archive (SRA) under the BioProject accession number PRJNA725120. Preprocessed count tables are available from `https://github.com/slhogle/consumer-competition`. Scripts reproducing all figures and steps in the data analysis are available from `https://github.com/slhogle/consumer-competition`.

**Words in Abstract:** 167

**Words in Main Text:** 5044

**Cited References in Main Text:** 53

**Figures in Main Text:** 5

**Keywords:** intraspecific variation | traits | coexistence | competition | predator | prey | community assembly | bacteria | nematodes | protists

This PDF file includes:
Supplementary Text
Supplementary Figures S1 to S4
Supplementary Tables S1 to S12

### Contents

**1 Supplementary Methods:** **4**

1.1 Study species source: 4

1.2 Experimental preparations: 4

1.3 Experimental sampling: 4

1.4 Bacterial trait measurements: 4

1.5 Consumer feeding efficiency experiments: 5

1.6 DNA extraction and sequencing: 5

1.7 DNA sequence analysis: 5

1.8 Reporting parameters from Bayesian models: 6

1.9 Bias estimates and calibration: 6

1.10 Statistical models of consumer and prey biomass: 6

1.11 Parameterization of the Lotka-Volterra competitive models: 7

1.12 Calculation of species diversity: 7

1.13 Calculation of diversity dissimilarity: 8

1.14 Identification of the sorting and equilibrium phase: 8

1.15 Comparisons of prey diversity across consumer treatments: 9

1.16 Joint species distribution models: 9

### List of Supplementary Figures

S1 **Bacterial prey relative abundance and diversity dissimilarities** 12

S2 **Identifying sorting and equilibrium phases from prey Shannon diversity** 13

S3 **Prey Shannon diversity in sorting and equilibrium phases** 14

S4 **Measured and predicted bacterial traits** 15

### 1 Supplementary Methods:

#### 1.1 Study species source:

The prey bacterial community consisted of 24 species (Table S1) from the University of Helsinki Culture Collection. The consumer species consisted of the ciliate *Tetrahymena thermophila* strain 1630/1U (CCAP), obtained from the Culture Collection of Algae and Protozoa at the Scottish Marine Institute (Oban, Scotland, United Kingdom) and the clonal hermaphrodite nematode *Caenorhabditis elegans* strain N2, obtained from the *Caenorhabditis* Genetics Center (University of Minnesota, Minneapolis, MN, USA).

#### 1.2 Experimental preparations:

Bacteria were revived from cryopreserved glycerol stocks (-80°C ) in protease peptone yeast (PPY) broth at 28°C with shaking for 24 hours. Individual colonies were picked from cultures plated on 50% PPY agar. Clones were then grown in 5% King's Broth (KB) until mid-log phase and cell densities (colony forming units) determined via plating on 50% PPY agar. Ciliates were revived from liquid nitrogen and cultured axenically at room temperature in PPY as described before[1, 2]. 50 ml of each revived ciliate culture was harvested by centrifugation (8 minutes, 3300 rpm, 4°C), then the cell pellet was resuspended in 25 ml M9 buffer. Ciliates were then counted by microscope using 40× magnification. Cryopreserved (-80°C) nematode stocks were revived following the standard protocol [3]. Briefly, frozen nematodes were spread on NGM-plates with *E. coli* OP50, then individuals transferred to fresh plates by "chunking" after eight days. Eggs were collected from 7-8 plates after six days by bleaching live worms. Developmentally synchronized nematodes were hatched in 125 ml M9 buffer with gentle rotation in tissue flasks overnight. Flasks were chilled on ice, 75 ml of buffer was manually decanted, and the remaining buffer and nematodes were centrifuged (3 mins, 1300 rpm, 8°C). Afterward, half the remaining buffer was decanted, and nematodes were counted directly by microscope.

#### 1.3 Experimental sampling:

One ml of each experimental treatment was sampled at each transfer, combined with 0.5 ml 85% glycerol, and frozen at -80 °C for later DNA extraction. Another 0.5 ml was sampled at each transfer for quantifying consumer density (ciliate: 1:3 vol/vol with 10% Lugol, nematode: 1:1 vol/vol with 20% Lugol). Nematode cell densities were measured directly under a microscope. Ciliate densities were measured using Fiji/ImageJ with the cell counter plugin[4].

#### 1.4 Bacterial trait measurements:

We measured innate predator defense, carrying capacity and growth rate for each bacterial strain as described in detail earlier[5]. Briefly, we revived each bacterial species in fresh medium (1% Kings Broth), acclimated them 24 hours, then pin replicated strains to fresh medium in 100 well plates for use with a Bioscreen C spectrophotometer (Growth Curves ABLtd, Helsinki, Finland). Plates were incubated at 28°C at constant shaking in the Bioscreen C and measurements were taken at 5 min intervals for 24 hours. After 48 hours, LTV ciliates were added to Bioscreen C wells to estimate biomass loss due to predation. For each bacterial species k we quantified growth potential using area under the logistic growth curve (AUC), and estimated innate defense of species k as $AUC_{k+consumer}/AUC_k$. We inferred maximum growth rate, the doubling time, the carrying capacity using growthcurveR[6]. We determined biofilm formation capacity for each using the crystal-violet method[7]. We determined bacterial growth on 31 different carbon substrates using BioLog EcoPlates (https://www.biolog.com).

#### 1.5 Consumer feeding efficiency experiments:

The 24 bacteria species, LTV ciliate, HTV ciliate, and nematode stocks were revived from cryopreservation as described above. Each of the bacterial species was grown as monocultures in 100 ml of Reasoner's 2A liquid medium[8] at room temperature for 96 hours with shaking at 50 RPM. Afterward, bacteria were harvested by centrifugation (13000 RPM, 10 minutes, 21°C), the supernatant was discarded, and cells were resuspended in 100 ml of M9 saline solution. The optical density was then adjusted to $OD_{600} = 0.5$ with additional M9. We added 5 ml of suspension from each bacterial species to 6 well culture plates and then added either $2.5 \times 10^3$ nematode individuals or $2.5 \times 10^4$ of HTV or LTV ciliate individuals. One well for each bacterial species was reserved as a no-consumer control. All conditions were assayed in biological triplicates. Plates were incubated at room temperature with shaking at 50 RPM. After 144 hours of incubation, we measured bacterial optical density ($OD_{600}$) and counted consumers as described above. We defined prey clearance as the difference between the $OD_{600}$ in each consumer treatment and the no consumer control. We converted optical density to cells $ml^{-1}$ using a conversion of $8 \times 10^6$ bacterial cells per 0.01 $OD_{600}$ unit. We defined prey clearance as the number of prey cells consumed per consumer individual after 144 hours.

#### 1.6 DNA extraction and sequencing:

Total DNA was extracted using the DNeasy Blood & Tissue 96-well extraction kit (Qiagen) according to the manufacturer's instructions. The V3-V4 region of the bacterial 16S rRNA was amplified using a primer set with Illumina sequencing adapters and multiplexing barcodes. Samples were sequenced on an Illumina MiSeq using v3 600 cycles reagent kit. The procedure is outlined in detail in earlier studies[9, 10].

#### 1.7 DNA sequence analysis:

Quality control and processing were performed using BBTools (version 38.61b (https://sourceforge.net/projects/bbmap/). 16S amplicon read pairs were processed using BBDuk to remove contaminants, trim reads that contained adapter sequence, and right quality trim reads where quality drops below Q10. BBMerge [11] was used to merge surviving read pairs into single amplicon sequences, and `msa.sh` and `cutprimers.sh` were used to remove any forward and reverse primer sequences. VSEARCH [12] was used to filter out amplicons with more than two expected errors (`-fastq_maxee 2`)[13] and excluded sequences outside of the 360-480 bp range. Quality controlled amplicon sequences were assigned to bacterial species using BBMap by mapping against a database of 30 full-length 16S rRNA sequences using the best possible mapping position. Only amplicons mapping unambiguously between position 341 to 805 in the aligned 16S rRNA region with at least a 95% mapping identity were retained and assigned to a species for subsequent analysis. Nearly all reads from each sample were perfectly mapped to a single species 16S rRNA sequence using these criteria, and the greatest number of unmapped reads was 0.1% of the total.

#### 1.8 Reporting parameters from Bayesian models:

We report parameter medians as an index of centrality and use the 95% Credible Interval (95% CI) as a measure of parameter uncertainty. We use the Probability of direction (Pd) as a measure of effect existence[14, 15]. Pd is correlated with the frequentist p-value (e.g., $Pd \leq 95\% \rightarrow P > 0.1$; $Pd > 95\% \rightarrow P < 0.1$; $Pd > 97\% \rightarrow P \approx 0.05$; $Pd > 99\% \rightarrow P \approx 0.01$; $Pd > 99.9\% \rightarrow P \leq 0.001$). We use the region of practical equivalence (ROPE) as a measure of effect size or significance that relates whether an estimated parameter reflects a non-negligible change (in terms of magnitude)[14, 15]. We report the percentage of the full posterior within the ROPE. Guidelines for interpreting the percent in ROPE: $> 99\%$ in ROPE $\rightarrow$ accept null; $> 97.5\%$ in ROPE $\rightarrow$ probably negligible; $\leq 97.5\%$ & $\geq 2.5\%$ in ROPE $\rightarrow$ undecided significance; $< 2.5\%$ in ROPE $\rightarrow$ probably significant; $< 1\%$ in ROPE $\rightarrow$ reject null.

For ease of interpretation, we highlight in red all Bayesian model terms where $Pd > 97\%$ which is equivalent to $P \lessapprox 0.05$.

#### 1.9 Bias estimates and calibration:

Because we knew the density at which each strain was initially inoculated (~$1 \times 10^4$ cfu ml$^{-1}$), we could quantitatively estimate the total bias associated with the combined steps of DNA extraction, PCR amplification, and sequencing[16]. Briefly, we first calculate a compositional error matrix between the observed and actual species compositions, then estimate bias as the compositional mean of the errors in the control samples of defined composition, and finally, we subtract this bias estimate from the observed relative abundances of each species to calibrate species relative abundances in experimental samples. This process was performed using the metacal v0.1.0 R package.

#### 1.10 Statistical models of consumer and prey biomass:

Ciliate density, nematode density, and bacterial optical density were modeled with hierarchical generalized additive models [17] implemented with the R package mgcv[18]. For the ciliate and nematode in $NC_{LTV}$ and $NC_{HTV}$ treatments, we replaced zero counts with simulated values drawn from the log normal distribution centered on the mean of the detection limit (100 ciliate ml$^{-1}$ and 5 nematodes ml$^{-1}$). This was done to improve model convergence and general performance. Data points in Fig. 2A are observations and do not include the zero-replaced values. In each model, the treatment (consumer x trait variability) combinations were modeled as parametric covariates. A global nonparametric smooth term was included for time, as well as a nonparametric treatment-specific time smooth. There was no detectable time-lagged autocorrelation so we omitted a residual correlation structure in the models. Basis dimensions ($k$) (i.e, the upper limit on degrees of freedom for each smooth) were chosen following the recommendations of Wood[18]. The response variable error distribution models were negative binomial with a log link function for ciliate and worm and Gaussian with a log-link for bacterial optical density. We obtained Estimated Marginal Means[19] and computed contrasts from each model using emmeans v1.5.0[20].

The bacterial population density was not significantly different between consumer species grown in monoculture or competing in co-culture. In the absence of consumers, bacterial density rapidly grew and stabilized on the fifth day. The presence of one or two consumer species significantly lowered prey biomass relative to no consumer condition. However, time-averaged bacterial biomass was independent of the consumer treatment (mean across consumer treatments, $OD_{600} = 0.22 \pm 0.06$), with marginally higher bacterial biomass in the HTV ciliate treatments and lower in the nematode-only treatment (Tables S3). In all consumer treatments, prey biomass decreased rapidly after experiment onset and was stable after five days with the exception of the nematode-only treatment.

The ciliate and nematode grew from starting densities of $10^4$ and 10 individuals ml$^{-1}$, respectively. In the absence of competition, the LTV ciliate, the HTV ciliate, and the nematode stabilized at $4.2 \pm 2.3 \times 10^4$, $5.0 \pm 2.3 \times 10^4$, and $6.0 \pm 2.5 \times 10^3$ individuals ml$^{-1}$, respectively, after an exponential growth period of about 17 days.

#### 1.11 Parameterization of the Lotka-Volterra competitive models:

We estimated differences in competitive ability and niche overlap by parameterizing phenomenological Lotka-Volterra competition models[21]. The Lotka-Volterra models make no specific assumptions about the mechanisms underlying species interactions so they can flexibly parameterized such that they approximate any combination of potential mechanisms. We used models taking the following form:

$$\frac{dN_i}{dt} = N_i(r_i - \alpha_{ii}N_i - \alpha_{ij}N_j)$$

Here $N_i$ is the abundance of the ciliate, $N_j$ is the abundance of the nematode, $t$ is time in days, $r_i$ is the per capita growth rate of the ciliate, $\alpha_{ii}$ is the linear effect of intraspecific competition (sometimes parameterized as carrying capacity, $K = \frac{1}{\alpha_{ii}}$) and $\alpha_{ij}$ is the linear effect of interspecific competition of the nematode on the ciliate.

We used gauseR v1.0[22] to **1)** determine time-lagged abundance for each species, **2)** estimate per capita growth rates, **3)** regress per captia growth on species abundances which allows parameterizing the Lotka-Volterra competition models, and **4)** simulated abundance dynamics and tuned Lotka-Volterra model parameters to match model output to observations. Using the empirically estimated Lotka-Volterra coefficients we estimated niche overlap simply as the geometric mean ratio of the intraspecific and interspecific coefficients:

$$\rho = \sqrt{\frac{\alpha_{ij}\alpha_{ji}}{\alpha_{ii}\alpha_{jj}}}$$

Where niche difference between the two species is just:

$$1 - \rho$$

Finally, we estimated the competitive ratio of the ciliate to the nematode as the ratio of intrinsic growth rates multiplied by the geometric mean ratio of each species competition sensitivity coefficients

$$\frac{k_i}{k_j} = \frac{r_i}{r_j} \times \sqrt{\frac{\alpha_{jj}\alpha_{ji}}{\alpha_{ii}\alpha_{ij}}}$$

#### 1.12 Calculation of species diversity:

We estimated Shannon diversity using DivNet[23], which provides population-level estimates of species diversity that account for the inherent compositional structure of microbiome data and species co-occurrence patterns/dependencies.

#### 1.13 Calculation of diversity dissimilarity:

We calculated a Shannon diversity dissimilarity index following Tuomisto[24, 25] and Jasinska[26] and references therein. Briefly, the dissimilarity among species composition in each treatment-sampling combination can be expressed as:

$$D' = \frac{M^q_{eff} - 1}{M - 1}$$

Where $M$ is the number of treatment-specific replicate populations under comparison (in this case four) and $M^q_{eff}$ is the effective number of distinct populations (i.e., the beta diversity [24]). Here $D' = 0$ with one effectively distinct population (i.e., all replicate populations contain the same species at the same frequencies) and $D' = 1$ if the effective population number equals the number of replicates (the replicates are completely distinct).

The parameter $q$ is a scaling factor for the Rényi diversity[27] (or the corresponding Hill number[28]), from which many common diversity metrics are special cases[29]. In the case of $q = 1$, Rényi diversity is the exponential of the Shannon diversity, and it can shown that $M^q_{eff}$ simply equals the exponential of the Jensen-Shannon divergence of the replicate populations[26]. Thus, for our study we define:

$$M^{q=1}_{eff} = \exp\left[\frac{1}{M} \sum_{population_p} \sum_{lineage_k} x^{(p)}_k \log\left(\frac{x^{(p)}_k}{\bar{x_k}}\right)\right]$$

where $x_k$ is the frequency of species $k$ in replicate sample $p$.

For reference, we estimated the higher range of $D'$ that we might expect if the 24 species trajectories from our experiments were as variable across replicates as they were across consumer treatment categories. We performed a simple permutational procedure randomly assigning species abundances across replicates and treatment categories and then calculating $D'$ for the permuted samples. Since the value $D'$ is bounded on the unit interval, we used Bayesian beta regression to test the difference between permuted $D'$ values and observed values. The observed $D'$ values (Fig. S1B) were close to 0 (highly similar replicates) and significantly lower than expected if cross-replicate variance equaled cross-treatment variance (Table S5).

#### 1.14 Identification of the sorting and equilibrium phase:

We used segmented regressions implemented in the MCP R package v0.3.0 [30] to identity sorting and equilibrium phases using Shannon diversity. We used default priors, ran three independent MCMC chains, and sampled both the prior and the posterior with 10000 post warmup samples for each chain. We assessed convergence using effective sample sizes (approaching $> 10000$ for each parameter)[31] and potential scale reduction factors ($\hat{R} < 1.05$ for all covariates)[32]. We fit three models with alternative assumptions and compared the predictive performance of all models using Leave-One-Out Cross-Validation (LOO-CV) implemented in the R package loo v2.3.1[33]. We compared models by relative changes in Estimated Log Predictive Density, selecting the model with the highest value. The final selected model is described in the results section. After defining the phase transition point, we partitioned sample points into the two phases and modeled Shannon diversity independently for each phase using generalized linear models. We included a fixed time effect, a treatment effect, and a time-treatment interaction effect for the sorting phase model and only a treatment effect for the equilibrium model. We compared model contrasts and compared slopes from estimated marginal means in the same way as for the consumer biomass models.

The segmented regression models allow for nonzero, experiment-wide fixed slopes in the first phase and the second phase. Including a treatment-dependent random effect for the change point significantly increased the out-of-sample predictions for the model [Bayesian LOO-CV, ELPD = -36.5, SE = 12.2]. Therefore, the consumer treatment had an important effect on the length of the sorting phase. We also found no support for a model allowing for a nonzero slope in the second phase [Bayesian LOO-CV, ELPD = -6.7, SE = 4.6], which suggests that diversity in the equilibrium phase was best modeled as a constant with different mean diversity across treatments. We selected a final model (Fig. 2C) with a fixed diversity decline in the sorting phase, a treatment-dependent sorting-phase length, and a fixed treatment-dependent mean diversity in the equilibrium phase. We estimated an experiment-wide onset of equilibrium of $12.6 \pm 3.1$ days (Fig. 2C).

#### 1.15 Comparisons of prey diversity across consumer treatments:

We modeled Shannon diversity using generalized linear models with an inverse Gaussian error distribution using an inverse quadratic link function. We used the emtrends package to estimate marginal means of linear trends for the statistical interaction between time and each treatment category.

#### 1.16 Joint species distribution models:

We used a joint species distribution model framework [34] to simultaneously evaluate prey species and community responses to different consumer competitive arrangements. In contrast to individual species models, JSDMs assume that species share general responses to the environment and to each other. Thus, JSDMs can model the magnitude and sign of individual species' responses to their environment as a function of shared species traits or phylogenetic relatedness. We used the Hierarchical Modelling of Species Communities (HMSC) R package[35, 36]. HMSC simultaneously estimates community- and species-level relationships to environmental covariates and how these relationships are influenced by species traits and phylogenetic relatedness among species. HMSC also infers residual species co-occurrence patterns (e.g., potential species interactions) not attributable to fixed covariates.

**Study design:** Joint species distribution models are powerful because they typically rely on the assumption of (generalized) linear relationships between covariates and species abundances. However, in our experiments, most species clearly exhibited nonlinear temporal trajectories. To partially account for this within the linear framework, we partitioned the dataset and constructed two independent models for the two distinct phases of species diversity (sorting and equilibrium) that we previously identified by segmented regression of the diversity data. Most species either increased or decreased linearly with time as the community sorted over the first 13 days. The rarest species had been effectively excluded after 17 days, after which the remaining species' abundances generally oscillated around a stable mean. In the sorting phase, we modeled all 24 prey species except for 0262 due to its universally low abundance. In the equilibrium phase, we excluded prey species 0097, 0262, 1842, 1988, 2159, 2160, 2164, 2443, 2494, 2792, 3031, 3237 because they were too rare to model reliably.

**Sampling unit and response variable:** We used individual temporally-resolved microcosm samples ($day \times replicate \times condition$) as the model sampling unit resulting in 192 discrete samples. We used log-transformed sequencing read counts as the response variable. We used a two-step hurdle model approach to account for zero inflation in the count data where we first fit a probit model for presence/absence response and then a Gaussian response model for regularized log-transformed species abundances conditional upon presence. We followed the general HMSC approaches from earlier studies using sequence count data with hurdle models [35–37].

**Predictors:** In all models, we included as fixed effects the sequencing depth (to account for different sequencing library sizes), the presence of LTV or HTV ciliate, and the presence of the nematode allowing for differing effects of the ciliate across strata of the nematode in models for both phases. In the sorting phase, we modeled both time and any treatment-specific differences over time ($time \times condition$ interaction) as fixed effects. In the equilibrium phase, time was only included as a random effect. For all models, we also included a temporally explicit random effect for days post-experiment initialization and a non-structured random effect corresponding to the level of the individually sampled microcosms to account for repeated measures observations from each microcosm. We tested three different combinations of fixed and random effects using both the standard and hurdle model approaches. The combinations included P1) a full model including fixed and random effects, species traits, and phylogenetic relationships P2) the same as P1 but without random effects, and P3) the same as P1 but without fixed effects but including the fixed effects for sequencing depth and time in the sorting phase. Our goal with these models was to determine to what extent of species variation is attributable to fixed effects (the consumer/evolution experimental treatment) versus residual between-species associations and latent effects.

**Traits and phylogeny:** We included a phylogenetic species correlation matrix to account for phylogenetic signal in the species responses. We inferred a Maximum Likelihood species phylogeny with multithreaded RAxML [38] using a general time-reversible evolution model and the GAMMA model of rate heterogeneity (GTRGAMMA). The phylogeny was based on a nucleotide alignment of the 16S rRNA sequence from MAFFT[39]. We also included measured bacterial traits to determine the influence of traits on the expected species responses to experimental covariates. Using genomic inference and experimental measurement (Fig. S4), we quantified 12 different bacterial traits reflecting innate defense against the ciliate, species growth rate, metabolic versatility, and various bacterial lifestyles. To avoid including co-linearities and trait redundancy in the models, we reduced the dimensionality of trait-space using Principal Components Analysis (PCA). We found that the majority of prey trait variability reduced to two principal components jointly explaining 60% of the variance and broadly reflecting a defense-growth rate trade-off (Fig. S4). Strains that were more defended against the LTV ciliate generally had lower maximum intrinsic growth rates and less metabolic versatility (measured as the number of different carbon compounds from a bioassay supporting growth). Biofilm production was strongly oriented along a third PC explaining 14% of the total variance. In the final JSDMs we included the traits for defense, growth rate, carbon use, and biofilm formation. To reduce the potential for overfitting trait data, we did not include all traits in the final model. We chose not to use principle coordinate loadings of the traits directly to ease interpretation. To remove potential correlations between the traits, we ensured that the four included trait vectors were generally orthogonal (aligned on distinct PC axes). The first three principal coordinates upon which these four traits aligned accounted for over 80% of the total trait variation for the full dataset.

**Model fitting:** We fitted models using four independent Markov Chain Monte Carlo (MCMC) chains, each chain consisting of $1.25 \times 10^5$ iterations with half discarded as burn-in and sampling every 250 iterations to yield 2000 total posterior samples over 0.5 million total iterations. The mixing of the chains was sufficient. The effective sample sizes approached 2000 for each species [31] and $\hat{R} < 1.05$ for all covariates[32].

**Post-processing:** We examined the explanatory and predictive powers of the probit models through species-specific Tjur's $R^2$ values[40]. The explanatory and predictive powers of the abundance models were measured by conventional $R^2$ (i.e., the proportion of the variance explained/predicted from the independent variables). Explanatory power was derived from model predictions with the data used to fit the model. To compute predictive power, we performed 5-fold cross-validation, setting aside four randomly partitioned folds for model training and one fold for testing/prediction. We partitioned the total explained variance among fixed and random effects. We used the 95% posterior probability level to determine whether species or trait responses to experimental consumer treatments existed. In practice, this requires that the 95% credible interval of the sampled posterior for a model coefficient is either positive or negative (not both).

### Supplementary Figures:

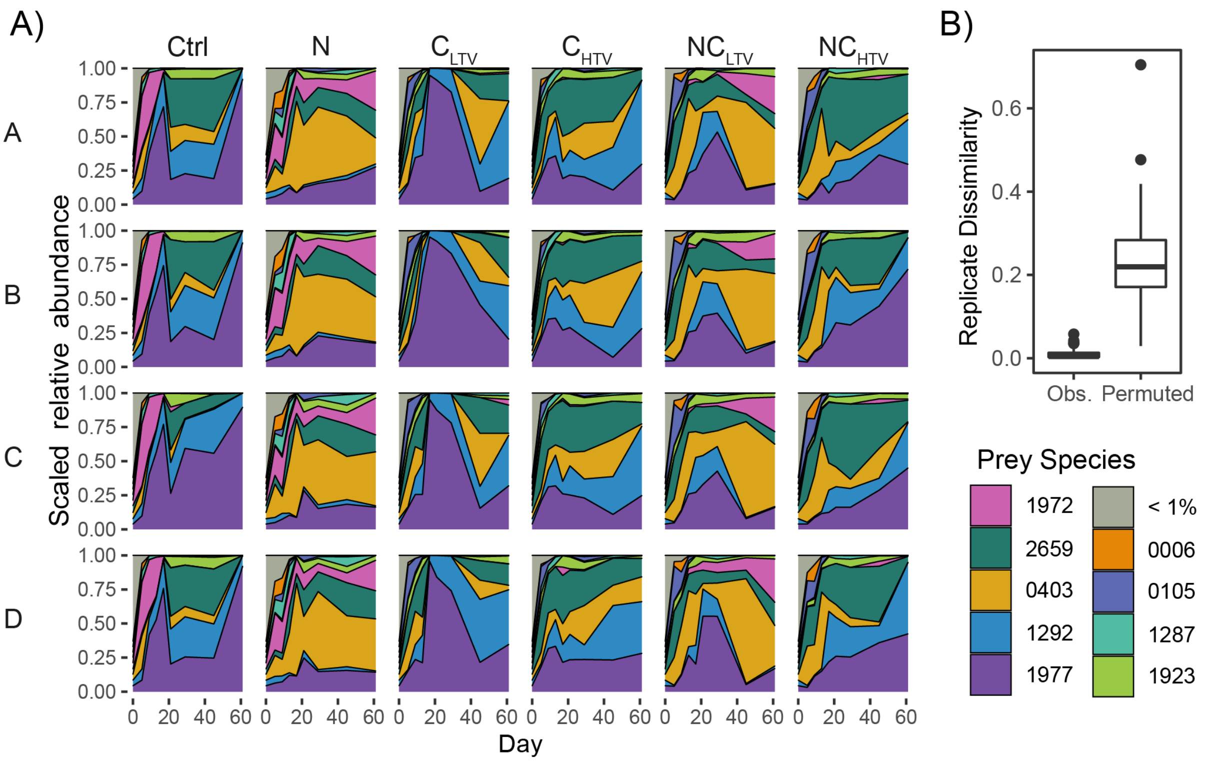

**Supplementary Figure S1. Bacterial prey relative abundance and diversity dissimilarities**
**A)** Relative abundance of the most abundant bacterial species in four biological replicates (rows, A-D) across 6 different experimental conditions (columns). The nine named strains constitute 99% of all reads across pooled experimental conditions. Ctrl - bacteria only; N - nematode; $C_{\mathrm{LTV}}$ - low trait diversity isogenic ciliate; $C_{\mathrm{HTV}}$ - high trait diversity ciliate; $NC_{\mathrm{LTV}}$ - nematode and low diversity ciliate; $NC_{\mathrm{HTV}}$ - nematode and high diversity ciliate. **B)** Shannon dissimilarity of lineages over replicate microcosms for each sampling time and consumer treatment. Obs - Observed Shannon dissimilarity; Permuted - Shannon dissimilarity for replicates permuted across treatment conditions (i.e., replicates as dissimilar as average dissimilarity between treatments.) $D' = 0$ with one effectively distinct population (i.e., all replicate populations contain the same species at the same frequencies) and $D' = 1$ if the effective population number equals the number of replicates (the replicates are completely distinct)

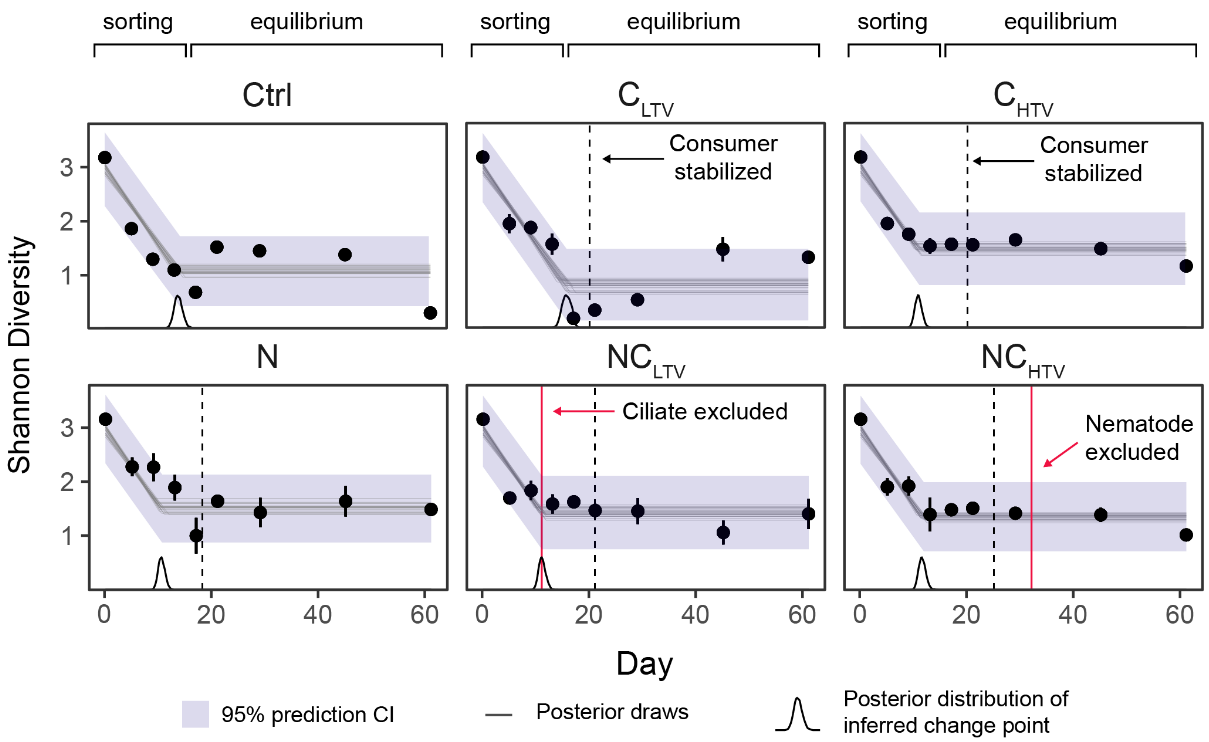

**Supplementary Figure S2. Identifying sorting and equilibrium phases from prey Shannon diversity** Shannon diversity of prey species over time. Points are observations with 95% confidence interval. The gray lines are 25 samples from the posterior of the model fit, the purple shading is the 95% credible interval of predictions from in-sample data, and the black distributions on the horizontal show the estimated equilibrium change point posterior distribution. Red lines in treatments $NC_{LTV}$ and $NC_{HTV}$ show when a competing consumer falls below starting density and is effectively excluded. Black dashed lines represent the time when consumer density stabilized. Ctrl - bacteria only, N - bacteria + nematode, $C_{LTV}$ - bacteria + low trait variance ciliate, $C_{HTV}$ - bacteria + high trait variance ciliate, $NC_{LTV}$ - bacteria + nematode + low trait variance ciliate, $NC_{HTV}$ - bacteria + nematode + high trait variance ciliate.

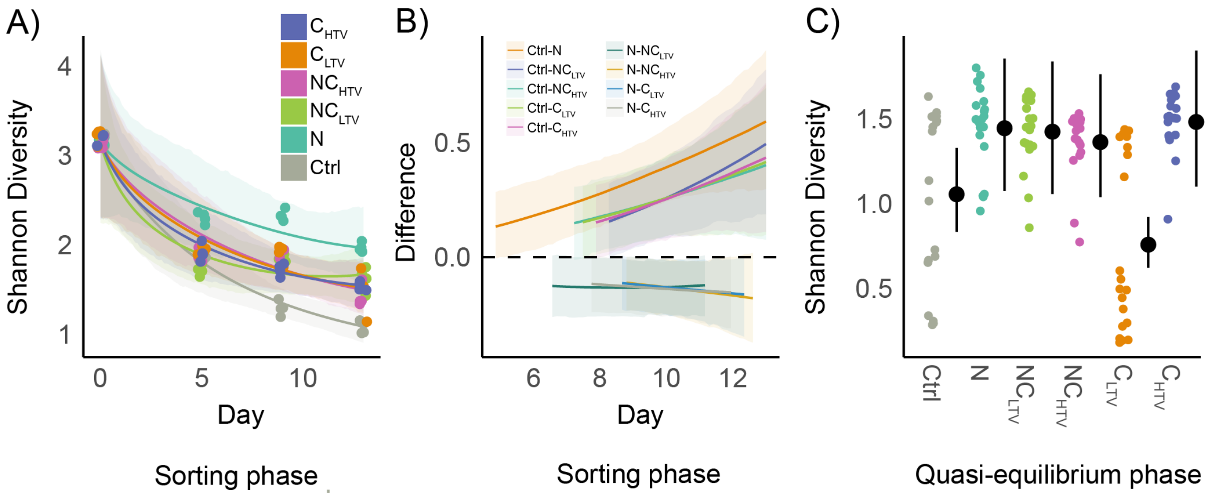

**Supplementary Figure S3. Prey Shannon diversity in sorting and equilibrium phases**

A) Observed Shannon diversity (points) and prediction (shaded lines) from a GLM modeling Shannon diversity as a function of time and consumer treatment in the sorting phase (days 0-16). Shaded regions are 95% Credible Intervals. B) Differences in Shannon diversity modeled from the GLM in A) at 100 equally spaced intervals of time. Contrasts are only shown if the difference between two conditions has $>$ 97% probability of direction. C) Observed Shannon diversity (points) and prediction (black point ranges) from a GLM modeling Shannon diversity as a function of consumer treatment only in the equilibrium phase (days 17-61). Ctrl - bacteria only, N - bacteria + nematode, $C_{LTV}$ - bacteria + low trait variance ciliate, $C_{HTV}$ - bacteria + high trait variance ciliate, $NC_{LTV}$ - bacteria + nematode + low trait variance ciliate, $NC_{HTV}$ - bacteria + nematode + high trait variance ciliate.

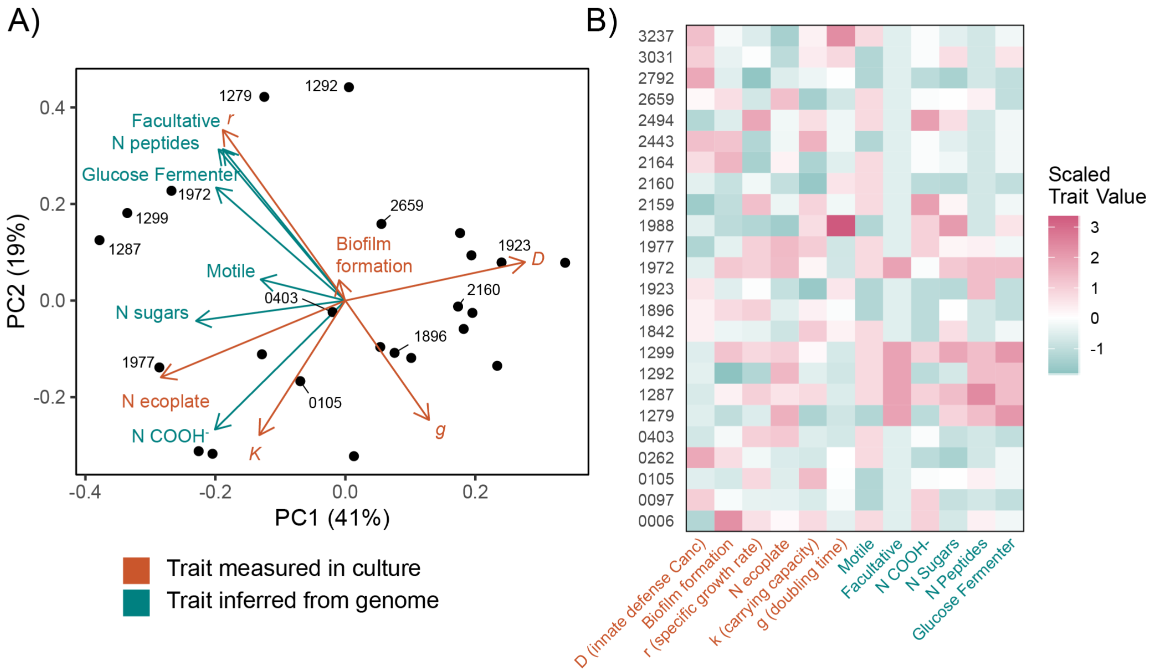

**Supplementary Figure S4. Measured and predicted bacterial traits**

The trait dataset consists of 12 discrete microbial traits, six of which were measured directly in culture (orange) and the six of which were predicted from whole genome sequences (green, see methods). A) Principal components analysis for the bacterial species with trait loadings. PC1 is dominated by the innate defense against the low trait variance, isogenic ciliate (D) and explains 41% of the trait variance in species. PC2 is largely dominated by the maximal specific growth rate (r) and explains 19% of total trait variance. B) Scaled traits plotted for each individual species. Scaled trait values are centered by the trait mean and normalized by the standard deviation. For traits measured in culture, D is defense against LTV ciliate; r is the maximum specific growth rate; N ecoplate is the number of carbon compounds used from the ecoplate bioassay; k is the maximum carrying capacity; and g is the doubling time. For traits predicted from genome sequences, Facultative is facultative anaerobic growth; N COOH- is the number of consumed carboxylic acid compounds; N Sugars is the number of consumed carbohydrate compounds; N Peptides is number of consumed peptides; and Glucose Fermenter indicates potential for growth via glucose fermentation.

### Supplementary Tables:

**Supplementary Table S1.** Bacterial species used in the experiment

| HAMBI collection strain ID | Species |
|---|---|
| 0006 | *Pseudomonas putida* |
| 0097 | *Acinetobacter johnsonii* |
| 0105 | *Agrobacterium tumefaciens* |
| 0262 | *Brevundimonas bullata* |
| 0403 | *Comamonas testosterone* |
| 1279 | *Hafnia alvei* |
| 1287 | *Citrobacter koseri* |
| 1292 | *Morganella morganii* |
| 1299 | *Kluyvera intermedia* |
| 1842 | *Sphingobium yanoikuyae* |
| 1896 | *Sphingobacterium spiritivorum* |
| 1923 | *Myroides odoratus* |
| 1972 | *Aeromonas caviae* |
| 1977 | *Pseudomonas chlororaphis* |
| 1987 | *Chitinophaga sancti* |
| 2159 | *Paraburkholderia caryophylli* |
| 2160 | *Bordetella avium* |
| 2164 | *Cupriavidus necator* |
| 2443 | *Paracoccus denitrificans* |
| 2494 | *Paraburkholderia kururiensis* |
| 2659 | *Stenotrophomonas maltophilia* |
| 2792 | *Moraxella canis* |
| 3031 | *Niabella yanshanensis* |
| 3237 | *Microvirga lotononidis* |

HAMBI = University of Helsinki Microbial Domain Biological Resource Centre

**Supplementary Table S2.** Ciliate density GAM

| **A) Model Summary** | | | | | |
|---|---|---|---|---|---|
| Parametric coefficients | Estimate | Std. Error | t-value | p-value | |
| (Intercept): $C_{LTV}$ | 10.5292 | 0.0493 | 213.6321 | $< 0.0001$ | |
| $C_{HTV}$ | 0.1707 | 0.0696 | 2.4537 | 0.0150 | |
| $NC_{LTV}$ | -3.9705 | 0.0701 | -56.6726 | $< 0.0001$ | |
| $NC_{HTV}$ | 0.0657 | 0.0694 | 0.9474 | 0.3446 | |
| Smooth terms | Est. df | Ref. df | F-value | p-value | |
| s(day) | 8.9657 | 10.0978 | 6.9961 | $< 0.0001$ | |
| s(day):$C_{LTV}$ | 9.8400 | 12.0000 | 4.8512 | 0.7522 | |
| s(day):$C_{HTV}$ | 8.1609 | 12.0000 | 326.8437 | 0.0140 | |
| s(day):$NC_{LTV}$ | 10.9902 | 12.0000 | 105.1107 | $< 0.0001$ | |
| s(day):$NC_{HTV}$ | 0.0000 | 12.0000 | 0.0000 | 0.0633 | |
| **B) Model Contrasts** | | | | | |
| Contrast | Ratio | Std. Error | t-ratio | p-value | Df |
| $C_{LTV}$ - $NC_{LTV}$ | 199.6925 | 42.6455 | 24.803 | <.0001 | 198.04 |
| $C_{HTV}$ - $NC_{LTV}$ | 334.3618 | 70.9935 | 27.374 | <.0001 | 198.04 |
| $NC_{LTV}$ - $NC_{HTV}$ | 0.0034 | 0.0007 | -29.020 | <.0001 | 198.04 |

**A)** GAM summary for ciliate density. Response error distributions were negative binomial with log-link. Smooth parameters were estimated using the fast mixed model approach via restricted maximum likelihood (fREML). Estimated coefficients are on the log scale. **B)** Pairwise model contrasts. For brevity only contrasts with $P \leq 0.05$ are included. P value adjustment: Bonferroni method for 6 tests. Tests are performed on the log scale

**Supplementary Table S3.** Bacterial density GAM

| **A) Model Summary** | | | | | |
|---|---|---|---|---|---|
| Parametric coefficients | Estimate | Std. Error | t-value | p-value | |
| (Intercept) | -0.3074 | 0.0076 | -40.4702 | < 0.0001 | |
| N | -1.2545 | 0.0322 | -38.9329 | < 0.0001 | |
| $C_{LTV}$ | -1.1987 | 0.0261 | -45.9421 | < 0.0001 | |
| $C_{HTV}$ | -1.0297 | 0.0236 | -43.6432 | < 0.0001 | |
| $NC_{LTV}$ | -1.2891 | 0.0288 | -44.7122 | < 0.0001 | |
| $NC_{HTV}$ | -1.0342 | 0.0226 | -45.8105 | < 0.0001 | |
| Smooth terms | Est. df | Ref. df | F-value | p-value | |
| s(day) | 1.0000 | 1.0000 | 0.0278 | 0.8678 | |
| s(day):Ctrl | 8.0301 | 11.0000 | 7.9842 | < 0.0001 | |
| s(day):N | 10.2130 | 11.0000 | 66.1397 | < 0.0001 | |
| s(day):$C_{LTV}$ | 6.2171 | 11.0000 | 1.7780 | 0.0005 | |
| s(day):$C_{HTV}$ | 10.3907 | 11.0000 | 7.4567 | < 0.0001 | |
| s(day):$NC_{LTV}$ | 6.7563 | 11.0000 | 3.1599 | < 0.0001 | |
| s(day):$NC_{HTV}$ | 9.1358 | 11.0000 | 4.1137 | < 0.0001 | |
| **B) Model Contrasts** | | | | | |
| Contrast | Ratio | Std. Error | t-ratio | p-value | Df |
| Ctrl - N | 4.9722 | 0.5344 | 14.923 | <.0001 | 278.26 |
| Ctrl - $C_{LTV}$ | 3.6096 | 0.2377 | 19.496 | <.0001 | 278.26 |
| Ctrl - $C_{HTV}$ | 2.5507 | 0.1484 | 16.096 | <.0001 | 278.26 |
| Ctrl - $NC_{LTV}$ | 4.3373 | 0.3425 | 18.579 | <.0001 | 278.26 |
| Ctrl - $NC_{HTV}$ | 2.7049 | 0.1602 | 16.803 | <.0001 | 278.26 |
| N - $C_{HTV}$ | 0.5130 | 0.0610 | -5.616 | <.0001 | 278.26 |
| N - $NC_{HTV}$ | 0.5440 | 0.0649 | -5.100 | <.0001 | 278.26 |
| $C_{LTV}$ - $C_{HTV}$ | 0.7066 | 0.0588 | -4.176 | 0.0006 | 278.26 |
| $C_{LTV}$ - $NC_{HTV}$ | 0.7493 | 0.0629 | -3.440 | 0.0101 | 278.26 |
| $C_{HTV}$ - $NC_{LTV}$ | 1.7005 | 0.1597 | 5.655 | <.0001 | 278.26 |
| $NC_{LTV}$ - $NC_{HTV}$ | 0.6236 | 0.0590 | -4.995 | <.0001 | 278.26 |

**A)** GAM summary for bacterial density. Response error distributions were Gaussian with log-link. Smooth parameters were estimated using the fast mixed model approach via restricted maximum likelihood (fREML). Estimated coefficients are on the log scale. **B)** Pairwise model contrasts. For brevity only contrasts with $P \leq 0.05$ are included. P value adjustment: bonferroni method for 15 tests. Tests are performed on the log scale.

**Supplementary Table S4.** Nematode density GAM

| **A) Model Summary** | | | | | |
|---|---|---|---|---|---|
| Parametric coefficients | Estimate | Std. Error | t-value | p-value | |
| (Intercept): N | 8.2782 | 0.0531 | 155.9162 | $< 0.0001$ | |
| $NC_{LTV}$ | -0.7325 | 0.0775 | -9.4461 | $< 0.0001$ | |
| $NC_{HTV}$ | -5.8017 | 0.0962 | -60.3108 | $< 0.0001$ | |
| Smooth terms | Est. df | Ref. df | F-value | p-value | |
| s(day) | 7.4909 | 7.8599 | 50.7313 | $< 0.0001$ | |
| s(day):N | 0.0000 | 8.0000 | 0.0000 | 0.0070 | |
| s(day):$NC_{LTV}$ | 7.0253 | 8.0000 | 33.1247 | $< 0.0001$ | |
| s(day):$NC_{HTV}$ | 6.6574 | 8.0000 | 51.3753 | $< 0.0001$ | |
| **B) Model Contrasts** | | | | | |
| Contrast | Ratio | Std. Error | t-ratio | p-value | Df |
| N - $NC_{HTV}$ | 743.9669 | 170.0777 | 28.923 | $<.0001$ | 155.83 |
| $NC_{LTV}$ - $NC_{HTV}$ | 775.2611 | 188.6798 | 27.337 | $<.0001$ | 155.83 |

**A)** GAM summary for nematode density. Response error distributions were negative binomial with log-link. Smooth parameters were estimated using the fast mixed model approach via restricted maximum likelihood (fREML). Estimated coefficients are on the log scale. **B)** Pairwise model contrasts. For brevity only contrasts with $P \leq 0.05$ are included. P value adjustment: Bonferroni method for 3 tests. Tests performed on the log scale.

**Supplementary Table S5.** Beta regression summary: $D'$

| Parameter | Median | 95% CI | Pd | ROPE | % in ROPE |
|---|---|---|---|---|---|
| (Intcpt) $D'$ rand. | -1.37 | [-1.54, -1.21] | 100 | ±0.18 | 0.00 |
| $D'$ obs. | -2.15 | [-2.48, -1.80] | 100 | ±0.18 | 0.00 |
| ($\varphi$) | 13.67 | [10.03, 18.02] | 100 | ±0.18 | 0.00 |

Coefficients are on the logit scale. The observed $D'$ is significantly lower than the random/permuted $D'$. $\varphi$ is the mean and precision parameter from beta regression.

**Supplementary Table S6.** Sorting phase alpha diversity regression

**A) GLM Summary**

| Parameter | Median | 95% CI | Pd | ROPE | % in ROPE |
|---|---|---|---|---|---|
| (Intercept) Ctrl | 0.46 | [0.36, 0.56] | 100.0 | ±0.10 | 0 |
| N | -0.27 | [-0.38, -0.16] | 100.0 | ±0.10 | 0 |
| $C_{LTV}$ | -0.18 | [-0.30, -0.06] | 99.84 | ±0.10 | 12.16 |
| $C_{HTV}$ | -0.17 | [-0.29, -0.05] | 99.81 | ±0.10 | 0.05 |
| $NC_{LTV}$ | -0.18 | [-0.31, -0.07] | 99.83 | ±0.10 | 9.39 |
| $NC_{HTV}$ | -0.17 | [-0.31, -0.06] | 99.86 | ±0.10 | 11.05 |
| $Day_1$ | 2.73 | [1.75, 3.79] | 100.0 | ±0.10 | 10.01 |
| $Day_1$:N | -2.16 | [-3.32, -1.08] | 100.0 | ±0.10 | 12.01 |
| $Day_1$:$C_{LTV}$ | -1.51 | [-2.79, -0.37] | 99.56 | ±0.10 | 0 |
| $Day_1$:$C_{HTV}$ | -1.57 | [-2.77, -0.39] | 99.59 | ±0.10 | 11.24 |
| $Day_1$:$NC_{LTV}$ | -1.78 | [-2.95, -0.61] | 99.88 | ±0.10 | 0.16 |
| $Day_1$:$NC_{HTV}$ | -1.47 | [-2.70, -0.29] | 99.14 | ±0.10 | 5.42 |
| $Day_2$ | 0.39 | [-0.49, 1.25] | 80.73 | ±0.10 | 0.59 |
| $Day_2$:N | -0.41 | [-1.43, 0.56] | 79.12 | ±0.10 | 12.17 |
| $Day_2$:$C_{LTV}$ | -0.40 | [-1.45, 0.69] | 76.59 | ±0.10 | 0.56 |
| $Day_2$:$C_{HTV}$ | -0.51 | [-1.56, 0.58] | 82.46 | ±0.10 | 11.30 |
| $Day_2$:$NC_{LTV}$ | -0.75 | [-1.81, 0.36] | 91.14 | ±0.10 | 0.47 |
| $Day_2$:$NC_{HTV}$ | -0.33 | [-1.39, 0.76] | 72.54 | ±0.10 | 10.14 |
| ($\lambda$) | 35.08 | [24.76, 46.37] | 100.00 | ±0.10 | 0 |

**B) Estimated linear slopes over treatment factors**

| Parameter | Median | 95% CI | Pd | ROPE | % in ROPE |
|---|---|---|---|---|---|
| Ctrl | -0.11 | [-0.19, -0.05] | 100.0 | ±0.002 | 0.00 |
| N | -0.07 | [-0.17, -0.00] | 99.08 | ±0.002 | 0.56 |
| $C_{LTV}$ | -0.08 | [-0.17, -0.02] | 99.85 | ±0.002 | 0.21 |
| $C_{HTV}$ | -0.07 | [-0.15, -0.02] | 99.99 | ±0.002 | 0.01 |
| $NC_{LTV}$ | -0.05 | [-0.11, -0.01] | 100.0 | ±0.002 | 0.03 |
| $NC_{HTV}$ | -0.09 | [-0.18, -0.03] | 99.99 | ±0.002 | 0.04 |

**C) Contrasts in linear slopes over treatment factors**

| Parameter | Contrast | 95% CI | Pd | ROPE | % in ROPE |
|---|---|---|---|---|---|
| Ctrl - $NC_{LTV}$ | -0.06 | [-0.16, 0.03] | 92.24 | ±0.002 | 1.94 |
| Ctrl - N | -0.04 | [-0.16, 0.08] | 78.03 | ±0.002 | 2.89 |
| Ctrl - $C_{HTV}$ | -0.04 | [-0.14, 0.07] | 79.41 | ±0.002 | 3.61 |
| $NC_{LTV}$ - $NC_{HTV}$ | 0.04 | [-0.05, 0.15] | 82.42 | ±0.002 | 3.55 |
| Ctrl - $C_{LTV}$ | -0.03 | [-0.13, 0.09] | 70.80 | ±0.002 | 4.09 |

**A)** Model summary. $\lambda$ is the reciprocal of the scale parameter from the inverse-Gaussian distribution. **B)** Estimated slopes of the linear relationship between alpha diversity and time across treatment levels. **C)** Contrasts in the slopes. Only the top 5 largest differences are included for brevity.

**Supplementary Table S7.** Equilibrium phase alpha diversity regression

**A) Model Summary**

| Parameter | Median | 95% CI | Pd | ROPE | % in ROPE |
|---|---|---|---|---|---|
| (Intcpt) Ctrl | 0.93 | [ 0.51, 1.36] | 100.0 | ±0.10 | 0.00 |
| N | -0.43 | [-0.96, 0.06] | 95.60 | ±0.10 | 7.71 |
| $C_{LTV}$ | 0.85 | [ 0.03, 1.66] | 95.19 | ±0.10 | 9.03 |
| $C_{HTV}$ | -0.45 | [-0.95, 0.05] | 92.33 | ±0.10 | 11.71 |
| NPanc | -0.41 | [-0.92, 0.09] | 98.10 | ±0.10 | 1.22 |
| NPevo | -0.36 | [-0.86, 0.16] | 96.53 | ±0.10 | 6.41 |
| ($\lambda$) | 3.64 | [ 2.74, 4.62] | 100.0 | ±0.10 | 0.00 |

**B) Model Contrasts**

| Parameter | Median | 95% CI | Pd | ROPE | % in ROPE |
|---|---|---|---|---|---|
| Ctrl - N | 0.43 | [-0.06, 0.96] | 95.60 | ±0.10 | 8.16 |
| Ctrl - $NC_{LTV}$ | 0.41 | [-0.09, 0.92] | 95.19 | ±0.10 | 8.72 |
| Ctrl - $NC_{HTV}$ | 0.36 | [-0.16, 0.86] | 92.33 | ±0.10 | 11.12 |
| Ctrl - $C_{LTV}$ | -0.85 | [-1.66, -0.03] | 98.10 | ±0.10 | 2.40 |
| Ctrl - $C_{HTV}$ | 0.45 | [-0.05, 0.95] | 96.53 | ±0.10 | 6.91 |
| N - $NC_{LTV}$ | -0.01 | [-0.40, 0.39] | 52.52 | ±0.10 | 39.05 |
| N - $NC_{HTV}$ | -0.06 | [-0.46, 0.35] | 62.08 | ±0.10 | 36.44 |
| N - $C_{LTV}$ | -1.27 | [-2.02, -0.54] | 100 | ±0.10 | 0.04 |
| N - $C_{HTV}$ | 0.03 | [-0.36, 0.39] | 55.46 | ±0.10 | 39.60 |
| $NC_{LTV}$ - $NC_{HTV}$ | -0.04 | [-0.44, 0.37] | 58.73 | ±0.10 | 37.30 |
| $NC_{LTV}$ - $C_{LTV}$ | -1.26 | [-2.02, -0.52] | 99.99 | ±0.10 | 0.04 |
| $NC_{LTV}$ - $C_{HTV}$ | 0.04 | [-0.33, 0.43] | 58.40 | ±0.10 | 39.31 |
| $NC_{HTV}$ - $C_{LTV}$ | -1.21 | [-1.98, -0.48] | 99.99 | ±0.10 | 0.04 |
| $NC_{HTV}$ - $C_{HTV}$ | 0.08 | [-0.30, 0.48] | 67.06 | ±0.10 | 35.99 |
| $C_{LTV}$ - $C_{HTV}$ | 1.29 | [ 0.56, 2.03] | 100.0 | ±0.10 | 0.01 |

**A)** GLM summary. $\lambda$ is the reciprocal of the scale parameter from the inverse-Gaussian distribution. **B)** Untransformed model contrast using inverse-Gaussian probability distribution

**Supplementary Table S8.** Ordination jump lengths

| **A) Model Summary** | | | | | |
|---|---|---|---|---|---|
| Parameter | Median | 95% CI | Pd | ROPE | % in ROPE |
| No consumer (Intcpt) | 4.10 | [3.03, 5.25] | 100 | ±0.10 | 0.00 |
| $C_{LTV}$ | -0.21 | [-1.79, 1.41] | 59.41 | ±0.10 | 9.57 |
| $C_{HTV}$ | 2.53 | [0.47, 4.61] | 99.38 | ±0.10 | 0.31 |
| N | 2.26 | [0.37, 4.43] | 98.78 | ±0.10 | 0.62 |
| $NC_{LTV}$ | 0.46 | [-1.23, 2.17] | 69.55 | ±0.10 | 7.42 |
| $NC_{HTV}$ | 2.06 | [0.04, 4.00] | 98.11 | ±0.10 | 1.04 |
| $\alpha$ | 1.88 | [1.51, 2.27] | 100 | ±0.10 | 0.00 |
| **B) Model Contrasts** | | | | | |
| Parameter | Contrast | 95% CI | Pd | ROPE | % in ROPE |
| No consumer - $C_{HTV}$ | -2.49 | [-4.57, -0.41] | 99.30 | 0.10 | 0.38 |
| No consumer - N | -2.23 | [-4.21, -0.14] | 98.81 | 0.10 | 0.68 |
| No consumer - $NC_{HTV}$ | -2.05 | [-4.03, -0.07] | 98.19 | 0.10 | 0.83 |
| $C_{LTV}$ - $C_{HTV}$ | -2.71 | [-4.78, -0.6] | 99.60 | 0.10 | 0.22 |
| $C_{LTV}$ - N | -2.44 | [-4.52, -0.51] | 99.30 | 0.10 | 0.29 |
| $C_{LTV}$ - $NC_{HTV}$ | -2.27 | [-4.26, -0.39] | 99.06 | 0.10 | 0.59 |

**A)** Model summary, **B)** contrasts between treatments, only significant differences are included. Coefficients and contrasts are on the inverse scale. $\alpha$ is the shape parameter from the Gamma distribution

**Supplementary Table S9.** Explanatory and predictive power of the JSDMs

| Phase | Model | Included effects | N | Explanatory $R^2$ | Predictive $R^2$ |
|---|---|---|---|---|---|
| Sorting | COP | Full | 23 | 0.90 ± 0.09 | 0.74 ± 0.27 |
| Sorting | COP | Fixed only | 23 | 0.88 ± 0.11 | |
| Sorting | COP | Random only | 23 | 0.40 ± 0.22 | |
| Equil | COP | Full | 12 | 0.58 ± 0.19 | 0.29 ± 0.24 |
| Equil | COP | Fixed only | 12 | 0.42 ± 0.21 | |
| Equil | COP | Random only | 12 | 0.14 ± 0.22 | |
| Sorting | PA | Full | 23 | 0.60 ± 0.20 | 0.53 ± 0.20 |
| Sorting | PA | Fixed only | 23 | 0.58 ± 0.20 | |
| Sorting | PA | Random only | 23 | 0.18 ± 0.13 | |
| Equil | PA | Full | 12 | 0.21 ± 0.10 | 0.14 ± 0.07 |
| Equil | PA | Fixed only | 12 | 0.02 ± 0.02 | |
| Equil | PA | Random only | 12 | 0.14 ± 0.07 | |

Phase indicates whether the model was for the sorting phase (days 0-13) or the equilibrium phase (days 17-61). Model type indicates where an abundance model conditional on presence (COP) or a presence/absence (PA) model was used. Included effects column shows if the model included both fixed and random effects (Full), fixed effects only, or random effects only. The random effects only model also included the non-focal fixed effects for sequencing depth and time in the sorting phase model. Explanatory/Predictive power is measured by the Tjur $R^2$ for the presence/absence models and by the standard $R^2$ for the COP abundance models. Values are the mean ± standard deviation over the number of species (N) included in each model. The explanatory measure of model fit (Explanatory $R^2$) is computed using the same data for model fitting. To account for potential overfitting, we also included a measure of predictive power (Predictive $R^2$) which is based on 5 fold cross validation of the subsampled dataset.

**Supplementary Table S10.** Proportion of variance explained by traits in the abundance JSDM

| Covariate | $R^2_\beta$ | $R^2_y$ | Experiment phase |
|---|---|---|---|
| time | 0.16 | 0.22 | sorting (days 0-13) |
| $C_{LTV}$ | 0.46 | 0.22 | sorting (days 0-13) |
| $C_{HTV}$ | 0.51 | 0.22 | sorting (days 0-13) |
| N | 0.22 | 0.22 | sorting (days 0-13) |
| time:$C_{LTV}$ | 0.55 | 0.22 | sorting (days 0-13) |
| time:$C_{HTV}$ | 0.52 | 0.22 | sorting (days 0-13) |
| time:N | 0.11 | 0.22 | sorting (days 0-13) |
| N:$C_{LTV}$ | 0.61 | 0.22 | sorting (days 0-13) |
| N:$C_{HTV}$ | 0.63 | 0.22 | sorting (days 0-13) |
| time:N:$C_{LTV}$ | 0.31 | 0.22 | sorting (days 0-13) |
| time:N:$C_{HTV}$ | 0.28 | 0.22 | sorting (days 0-13) |
| $C_{LTV}$ | 0.51 | 0.44 | equilibrium (days 17-61) |
| $C_{HTV}$ | 0.62 | 0.44 | equilibrium (days 17-61) |
| N | 0.63 | 0.44 | equilibrium (days 17-61) |
| N:$C_{LTV}$ | 0.77 | 0.44 | equilibrium (days 17-61) |
| N:$C_{HTV}$ | 0.80 | 0.44 | equilibrium (days 17-61) |

Covariate is the term or interaction between terms from the JSDM. $R^2_\beta$ is the proportion of variance that traits explain for the response of individual species to each covariate or covariate interaction. $R^2_y$ is the proportion of variance that traits explain for different species abundances. Experiment phase denotes the phase of the experiment modeled by the JSDM. In the sorting phase, the JSDM included a fixed effect for the difference over time, and any condition-specific differences over time as well as a temporal random effect. In the equilibrium JSDM time is only included as a random effect. $C_{LTV}$, ancestral ciliate present; $C_{HTV}$, evolved ciliate present; N, nematode present. The traits included in the JSDM are defense against $C_{LTV}$, the number of carbon sources the strain can use, the maximum specific growth rate, and the relative fluorescence intensity of a biofilm stain under exponential growth.

**Supplementary Table S11.** Prey clearance regression results

| **A) Model Summary** | | | | | |
|---|---|---|---|---|---|
| Parameter | Median | 95% CI | Pd | ROPE | % in ROPE |
| (Intcpt) $C_{LTV}$ | 7.67 | [7.37, 8.00] | 100 | ±0.17 | 0.00 |
| $C_{HTV}$ | 0.74 | [0.26, 1.21] | 99.9 | ±0.17 | 0.00 |
| Nematode | 3.36 | [2.91, 3.84] | 100 | ±0.17 | 0.00 |
| **B) Model Contrasts** | | | | | |
| Parameter | Median | 95% CI | Pd | ROPE | % in ROPE |
| $C_{LTV}$ - $C_{HTV}$ | -0.74 | [-1.21, -0.26] | 99.9 | ±0.10 | 0.00 |
| $C_{LTV}$ - N | -3.36 | [-3.84, -2.91] | 100 | ±0.10 | 0.00 |
| $C_{HTV}$ - N | -2.63 | [-3.10, -2.15] | 100 | ±0.10 | 0.00 |

**A)** GLM summary with Gaussian probability distribution. **B)** Model contrasts. Response variable and contrasts are log transformed

**Supplementary Table S12.** Prey specificity regression results

**A) Model Summary**

| Parameter | Median | 95% CI | Pd | ROPE | % in ROPE |
|---|---|---|---|---|---|
| (Intercept) | 0.03 | [0.01, 0.06] | 99.88 | ±0.01 | 1.68 |
| $C_{LTV}$ | 0.66 | [0.55, 0.77] | 100.0 | ±0.01 | 0.00 |
| $C_{HTV}$ | 0.04 | [0.01, 0.07] | 99.28 | ±0.01 | 4.71 |
| N | 0.05 | [0.02, 0.08] | 99.85 | ±0.01 | 0.00 |
| $C_{LTV}$:$C_{HTV}$ | -0.38 | [-0.54, -0.22] | 100.0 | ±0.01 | 0.00 |
| $C_{LTV}$:N | -0.57 | [-0.73, -0.41] | 100.0 | ±0.01 | 0.00 |

**B) Estimated slopes over treatment factors**

| Parameter | Median | 95% CI | Pd | ROPE | % in ROPE |
|---|---|---|---|---|---|
| $C_{LTV}$ | 0.66 | [0.55, 0.77] | 100.00 | 0.02 | 0.00 |
| $C_{HTV}$ | 0.29 | [0.18, 0.4] | 100.00 | 0.02 | 0.00 |
| N | 0.09 | [-0.02, 0.21] | 93.90 | 0.02 | 7.95 |

**C) Contrasts in slopes over treatment factors**

| Parameter | Contrast | 95% CI | Pd | ROPE | % in ROPE |
|---|---|---|---|---|---|
| $C_{LTV}$ - $C_{HTV}$ | 0.38 | [0.22, 0.54] | 100.00 | 0.02 | 0.00 |
| $C_{LTV}$ - N | 0.57 | [0.41, 0.73] | 100.00 | 0.02 | 0.00 |
| $C_{HTV}$ - N | 0.19 | [0.02, 0.34] | 99.12 | 0.02 | 1.25 |

**D) Hypothesis tests: Slope = 1**

| Hypothesis | Evidence ratio | Posterior Probability | Significant |
|---|---|---|---|
| $Slope_{C_{LTV}} > 1$ | 0.0 | 0.0 | |
| $Slope_{C_{LTV}} < 1$ | 570 | 1.0 | * |
| $Slope_{C_{HTV}} > 1$ | 0.0 | 0.0 | |
| $Slope_{C_{HTV}} < 1$ | $\infty$ | 1.0 | * |
| $Slope_{N} > 1$ | 0.0 | 0.0 | |
| $Slope_{N} < 1$ | $\infty$ | 1.0 | * |

**A)** Model summary, **B)** Estimated slopes of relationship between species relative abundance in the presence of a single consumer vs the absence of a single consumer, **C)** Contrasts in the slopes, **D)** Hypothesis test for the relationships between prey species abundance in the presence and absence of consumers. $\beta$ is the slope of the relationship. *: For one-sided hypotheses, the posterior probability exceeds 95%; for two-sided hypotheses, the value tested against lies outside the 95%-CI. Posterior probabilities of point hypotheses assume equal prior probabilities.
